# Adipose tissue eQTL meta-analysis reveals the contribution of allelic heterogeneity to gene expression regulation and cardiometabolic traits

**DOI:** 10.1101/2023.10.26.563798

**Authors:** Sarah M. Brotman, Julia S. El-Sayed Moustafa, Li Guan, K. Alaine Broadaway, Dongmeng Wang, Anne U. Jackson, Ryan Welch, Kevin W. Currin, Max Tomlinson, Swarooparani Vadlamudi, Heather M. Stringham, Amy L. Roberts, Timo A. Lakka, Anniina Oravilahti, Lilian Fernandes Silva, Narisu Narisu, Michael R. Erdos, Tingfen Yan, Lori L. Bonnycastle, Chelsea K. Raulerson, Yasrab Raza, Xinyu Yan, Stephen C.J. Parker, Johanna Kuusisto, Päivi Pajukanta, Jaakko Tuomilehto, Francis S. Collins, Michael Boehnke, Michael I. Love, Heikki A. Koistinen, Markku Laakso, Karen L. Mohlke, Kerrin S. Small, Laura J. Scott

## Abstract

Complete characterization of the genetic effects on gene expression is needed to elucidate tissue biology and the etiology of complex traits. Here, we analyzed 2,344 subcutaneous adipose tissue samples and identified 34K conditionally distinct expression quantitative trait locus (eQTL) signals in 18K genes. Over half of eQTL genes exhibited at least two eQTL signals. Compared to primary signals, non-primary signals had lower effect sizes, lower minor allele frequencies, and less promoter enrichment; they corresponded to genes with higher heritability and higher tolerance for loss of function. Colocalization of eQTL with conditionally distinct genome-wide association study signals for 28 cardiometabolic traits identified 3,605 eQTL signals for 1,861 genes. Inclusion of non-primary eQTL signals increased colocalized signals by 46%. Among 30 genes with ≥2 pairs of colocalized signals, 21 showed a mediating gene dosage effect on the trait. Thus, expanded eQTL identification reveals more mechanisms underlying complex traits and improves understanding of the complexity of gene expression regulation.

---

Genetic regulation of gene expression influences the etiology of complex traits.^1–3^ Many genome-wide association study (GWAS) signals are located in non-coding regions and lack obvious candidate genes or mechanisms.^2,4^ Integrating trait and disease GWAS signals with expression quantitative trait locus (eQTL) signals has identified candidate genes and their directions of effect relative to disease risk at thousands of loci^1,3–10^. However, most reported eQTL studies either have not explored or have had limited power to observe the complexities of genetic regulation beyond a single eQTL for each gene. Larger eQTL studies with greater power are needed to better understand the genetic architecture of gene expression and its impact on complex traits.

Both GWAS and eQTL loci exhibit allelic heterogeneity,^1,11–13^ and the detection of multiple association signals within a locus can reveal complex regulatory mechanisms.^14,15^ Simultaneous analysis of multiple signals associated with gene expression and complex traits in large sample sizes has the potential to identify more shared signals than previously described or predicted.^16,17^ One method to detect allelic heterogeneity in eQTLs is to identify conditionally distinct signals associated with expression of the same gene.^1,6–9,11–13,15,18^ Allelic heterogeneity is identified more frequently in eQTL studies with larger sample sizes,^15,18^ and the relatively modest sample sizes in most eQTL studies have resulted in limited power to detect more than one signal per gene. eQTL meta-analyses enable larger sample sizes, but few eQTL meta-analysis studies have identified non-primary signals (secondary, tertiary, quaternary, etc.).^15,18^ Identifying non-primary signals with individual-level data from multiple eQTL studies can be tedious,^19^ however methods exist to detect conditionally distinct signals with both summary statistics and individual-level data.^20^ ^21^

Although many eQTL are shared across tissues,^1,22,23^ some are tissue-specific,^2,24^ motivating studies in disease-relevant tissues. Adipose tissue is intrinsically linked to cardiometabolic diseases such as obesity and type 2 diabetes, plays a role in the management of dyslipidemia, and is a contributing factor in insulin resistance and metabolic disease pathogenesis.^25–27^ Additionally, subcutaneous adipose tissue is relatively accessible from research volunteers, in contrast to other tissues relevant for the pathophysiology of cardiometabolic diseases, such as visceral adipose, heart and liver, that are primarily obtained from disease cohorts or deceased individuals. Several subcutaneous adipose eQTL studies of relatively healthy individuals have been conducted with sample sizes up to 722 individuals^1,6,28–30^, but these studies have not been analyzed together.

Here, we introduce AdipoExpress, an eQTL meta-analysis of five studies, two of which have not been reported previously, with a total of 2,344 subcutaneous adipose tissue samples. We provide a widely applicable approach to effectively identify conditionally distinct eQTL signals across multiple studies and we illustrated the genetic and genomic characteristics of the eQTL and their corresponding genes. We then carried out colocalization analysis of distinct adipose eQTL signals with distinct GWAS signals from 28 cardiometabolic traits and detected thousands of shared signals. For sets of eQTL signals that colocalized with sets of GWAS signals for the same trait, we used Mendelian randomization to quantify gene dosage effects on traits. This expanded discovery of eQTL enabled us to identify new putative risk genes and mechanisms for cardiometabolic traits. The full marginal and conditional eQTL summary statistics are publicly available (see data availability), enabling further integration with additional GWAS and molecular QTL studies.

## Results

### eQTL meta-analysis gene and signal discovery

We performed a subcutaneous adipose tissue stepwise eQTL meta-analysis of conditionally distinct signals. We implemented a forward and backward selection model from five studies consisting of up to 2,344 individuals using 29,254 genes and 6.4 million variants with minor allele frequency (MAF) of ≥ 0.01 across autosomes and the X chromosome (**Table 1; Tables S1-S2;** **Figure 1**, **Figure S1, Figure S2**). Analyzing all genes tested in at least two studies, we identified 18,476 eQTL genes and 34,774 eQTL signals (*P* ≤ 1e-6) (**Table 1; Table S3**), which is >1.6-fold more eQTL genes and 2.3-fold more signals than any of the individual studies (**Figure 1A-B**). Each gene in the meta-analysis had an average of 1.9 eQTL signals, and 51% of the genes had at least two signals, compared to the maximum 27% in any individual study (**Figure 1**). Among the 34,774 eQTL signals, 47% would have been missed if we had only identified primary eQTL signals. Almost all study participants (2,256/2,344) were of European ancestry, and a meta-analysis of these individuals identified 18,345 eQTL genes and 34,216 signals (**Table 1; Table S4**); 98% of eQTL genes and 87% of eQTL signals were shared between the meta-analyses. As downstream colocalization analyses depend on genetic similarity between GWAS signals of primarily European ancestry individuals and the eQTL samples, subsequent analyses included only participants of European ancestry.

**Table 1:**
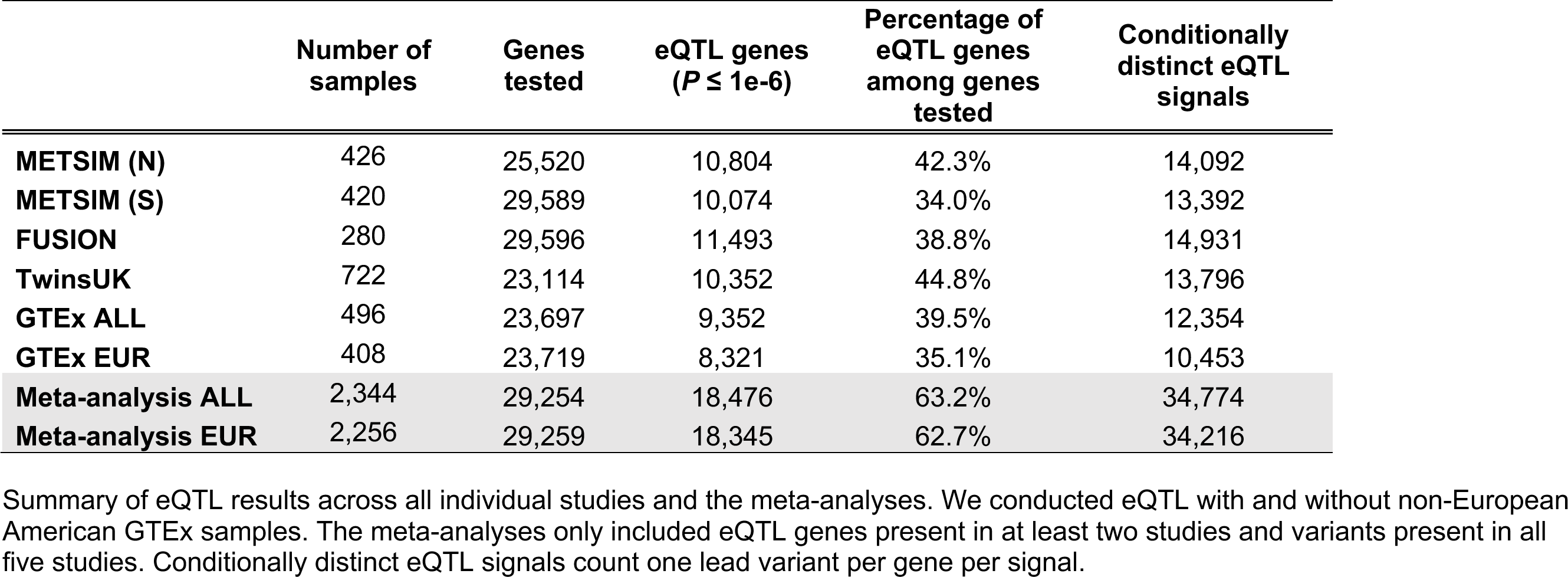
Overall discovery of adipose eQTL.

**Figure 1.**
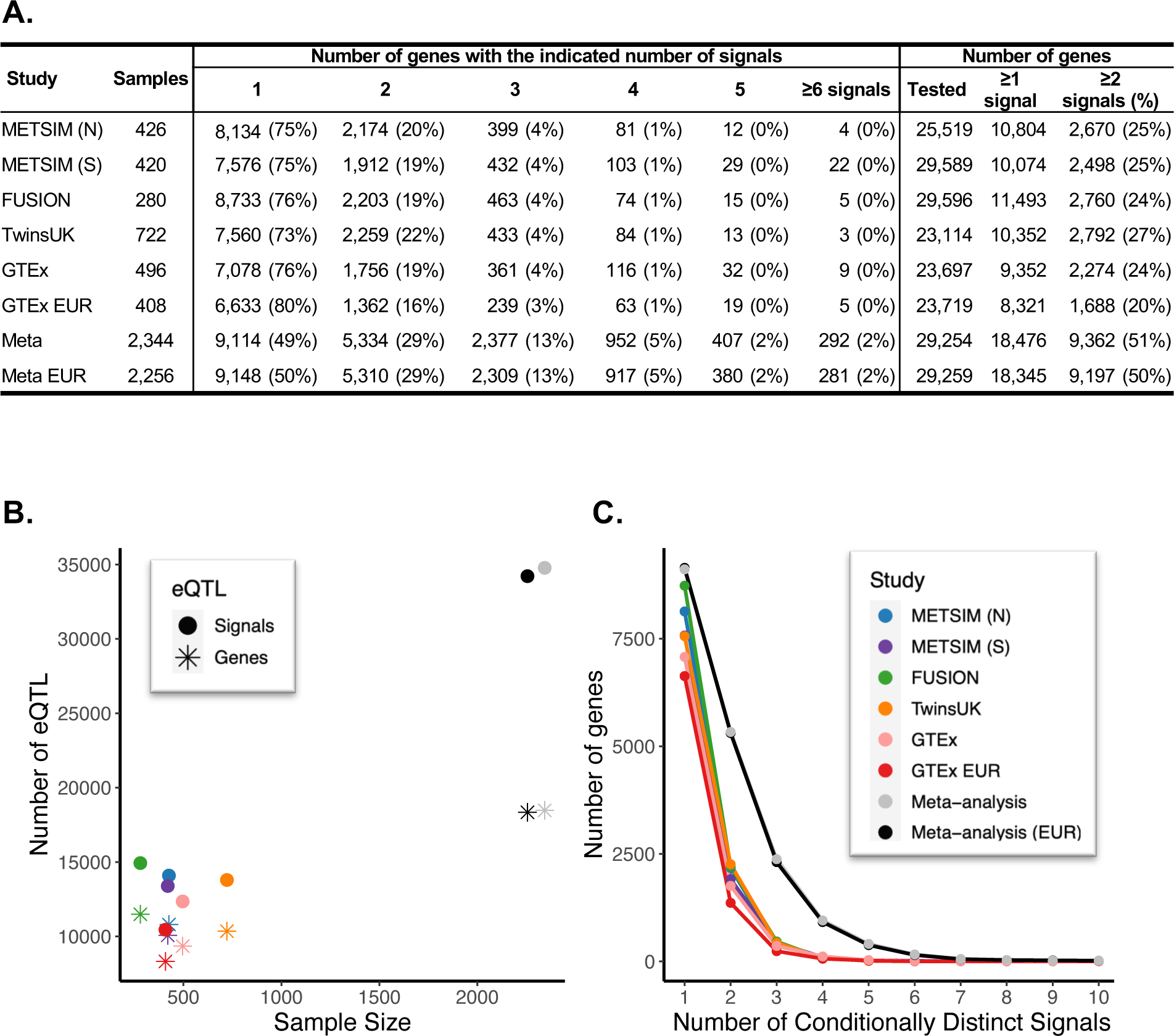
Conditionally distinct signals in adipose eQTL studies. A. Number of genes with 1 to 10 eQTL signals (*P* ≤ 1e-6) identified in each study and the meta-analyses. ‘≥1 signal’ column indicates the number of genes with at least one significant eQTL signal, ‘≥2 signals’ indicates the number of genes with two or more eQTL signals, and the percentage of genes with an eQTL that have two or more eQTL signals is in parentheses. B. The numbers of genes identified with an eQTL in each study are represented by filled circles, and the numbers of eQTL signals are represented by asterisks. Studies are shown by color: blue, METSIM (N); purple, METSIM (S); green, FUSION; orange, TwinsUK; pink, GTEx all populations; red, GTEx EUR; gray, meta-analysis with GTEx all populations; and black, meta-analysis with GTEx EUR. C. The number of genes with 1 through 10 eQTL signals detected in each study.

Due to the role of adipose tissue in GWAS traits with substantial sex differences^31^, we also conducted sex-stratified stepwise conditional eQTL meta-analyses using 270 female and 418 male individuals from the GTEx and FUSION studies that contained individuals of both sexes. We detected 8,473 eQTL genes and 10,510 eQTL signals in males and 6,834 eQTL genes and 8,035 eQTL signals in females (**Tables S5-S6**). Altogether, 45% of male eQTL signals are shared with female signals and 59% of female eQTL signals are shared with the male signals (LD r^2^ ≥ 0.8). The male and female marginal eQTL signals showed highly correlated effect sizes (Pearson r^2^ = 0.93) (**Figure S3**). Larger studies are needed to detect sex differences among adipose eQTL.

To relate eQTL discovery in adipose to a more accessible tissue, we compared the adipose eQTL signals to blood eQTL signals from the much larger eQTLGen^32^ study (n = 31,684). The studies had several differences in design (**Table S7**), including that eQTLGen reported only primary eQTL signals. Of the 18,345 primary adipose eQTL signals, 38% were potentially the same signal in blood (r^2^≥0.2), 29% corresponded to a gene not tested in blood, and 33% had an eQTL in blood that was not in LD (r^2^<0.2) with the adipose eQTL signal (**Figure S4; Table S8**). Of the 15,871 non-primary adipose eQTL signals, 21% were potentially the same signal in blood (r^2^≥0.2), 23% correspond to a gene not tested in blood, and 55% had an eQTL in blood that was not in LD (r ^2^<0.2) with the adipose eQTL signal (**Figure S4; Table S8**).Thus, even with a 10-fold smaller sample size in adipose than in blood, 62% of adipose eQTL were not detected as primary blood eQTL. A stepwise conditional analysis of eQTL signals in blood would likely detect additional signals shared across tissues.

### Characteristics of eQTL signals

Many eQTL studies only identify primary eQTL signals, and non-primary signals remain poorly characterized. Therefore, we compared characteristics of eQTL signals based on the order in which they were discovered in the stepwise conditional analysis. This order may depend on multiple factors, including effect sizes, minor allele frequencies, and cell-type composition, and can differ across studies. For example, at the *GLYCTK* gene, which encodes an enzyme involved in serine degradation and fructose metabolism, the meta-analysis identified two signals (signal 1 = chr3:52,273,421, rs610060; signal 2 = chr3:52,276,901, rs11711914; LD r^2^=0.14), while conditional analysis in the individual studies each only detected one significant signal (**Figure 2**). The individual studies identified different signals as significant: the studies with Finns identified signal 1 while the studies with non-Finnish Europeans identified signal 2 (**Figure 2**).

**Figure 2.**
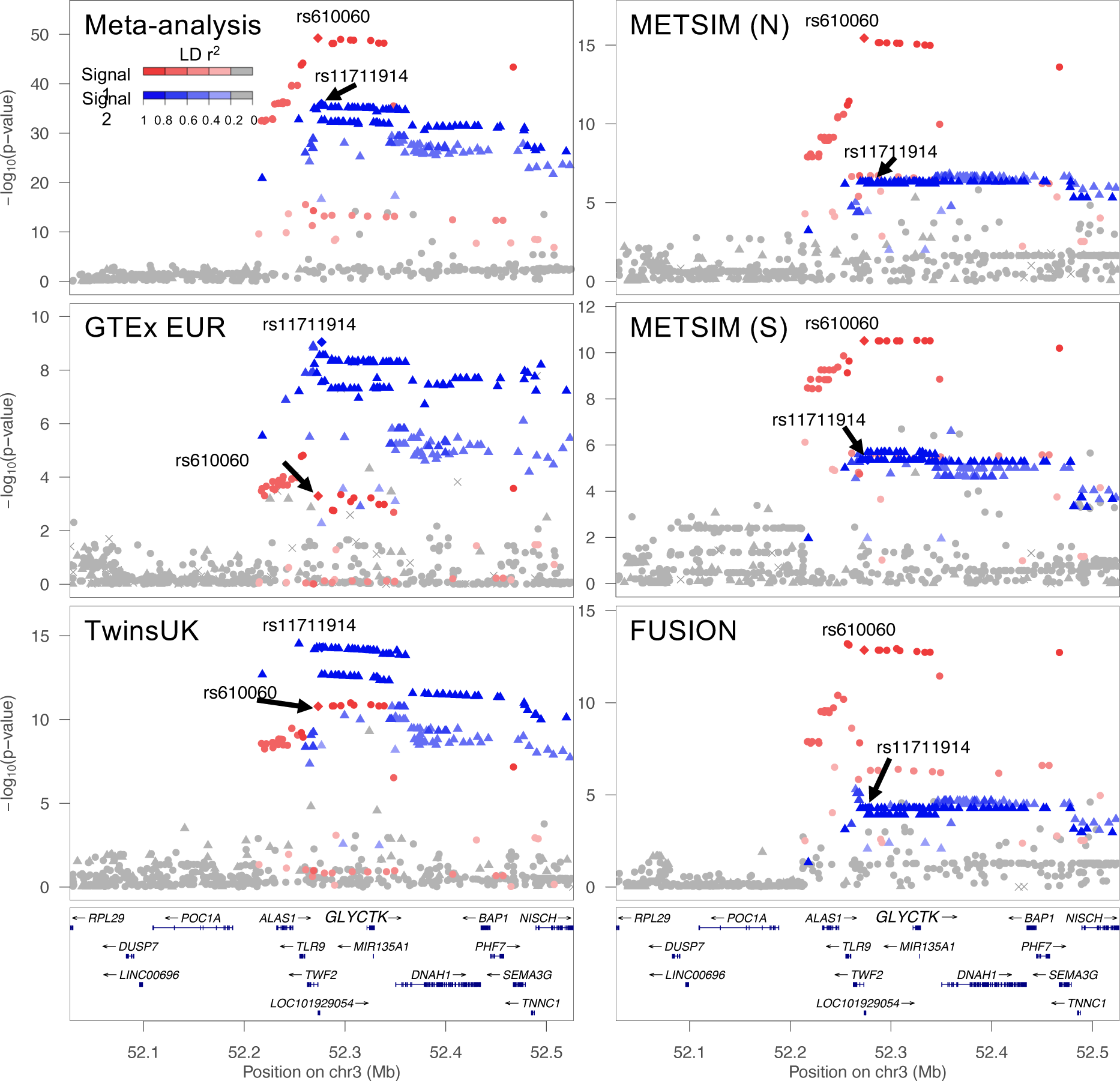
*GLYCTK* eQTL signals identified in each study and the meta-analysis. LocusZoom plots of the marginal *GLYCTK* eQTL for the meta-analysis and each individual study. The x-axes show position on chromosome 3 and y-axes show eQTL -log_10_ *P*-value. The lead variant of the 1^st^ signal (chr3:52,273,421, rs610060) in the meta-analysis is represented by a red diamond in all plots, and the lead variant of the second signal (chr3:52,276,901, rs11711914) in the meta-analysis is represented by a blue diamond in all plots. The red circles represent variants in stronger LD with the lead variant of the 1^st^ signal while the blue triangles represent variants in stronger LD with the lead variant of the second signal. Shading indicates LD r^2^ as shown in the legend. Although each study has both signals colored, only one signal was significant in the conditional eQTL analysis for each of the individual studies (*P* ≤ 1e-6).

Additionally, the lead variant allele frequencies differed between these populations, suggesting the difference in signal detection may be influenced by population (**Table S9**). Similarly, at the well-characterized *ADIPOQ* gene, the meta-analysis identified two signals in moderate pairwise LD (signal 1 = chr3:186,574,282, rs35469083; signal 2 = chr3:186,551,888, rs143257534; LD r^2^ = 0.35), while conditional analysis in the individual studies detected different single signals (**Figure S5; Table S10**). These examples show one way that the meta-analysis eQTL signals are more comprehensive than the signals detected by individual studies.

We compared primary and non-primary eQTL signals detected in the stepwise conditional analysis with respect to effect size, MAF, and distance to gene transcription start site (TSS). Effect sizes were typically lower for signals identified later; among the 661 genes with at least five eQTL signals, the median absolute value of the effect size for 1^st^ signals was twice as large as for 5^th^ signals (0.4 vs 0.2, *P* < 2e-16) (**Figure 3A**). In addition, MAF was typically lower for signals identified later; among genes with at least five signals, the median MAFs for 1^st^ and 5^th^ signals were 0.25 and 0.11, respectively (*P* < 2e-16) (**Figure 3B**). Finally, the distance from the lead eQTL variant to gene TSS became larger for signals identified later, indicating that the signals closest to a gene TSS tend to be discovered first. Among genes with five or more signals, the median distance to gene TSS was 26.4 kb for 1^st^ signals and 76.4 kb for 5^th^ signals (*P* < 2e-16) (**Figure 3C**). For all three characteristics, the same trends were observed for genes with two, three, or four signals (**Figure S6**). Thus, primary adipose eQTL signals had larger effect sizes, were discovered with more common variants, and the variants were closer to the TSS than subsequent signals.

**Figure 3.**
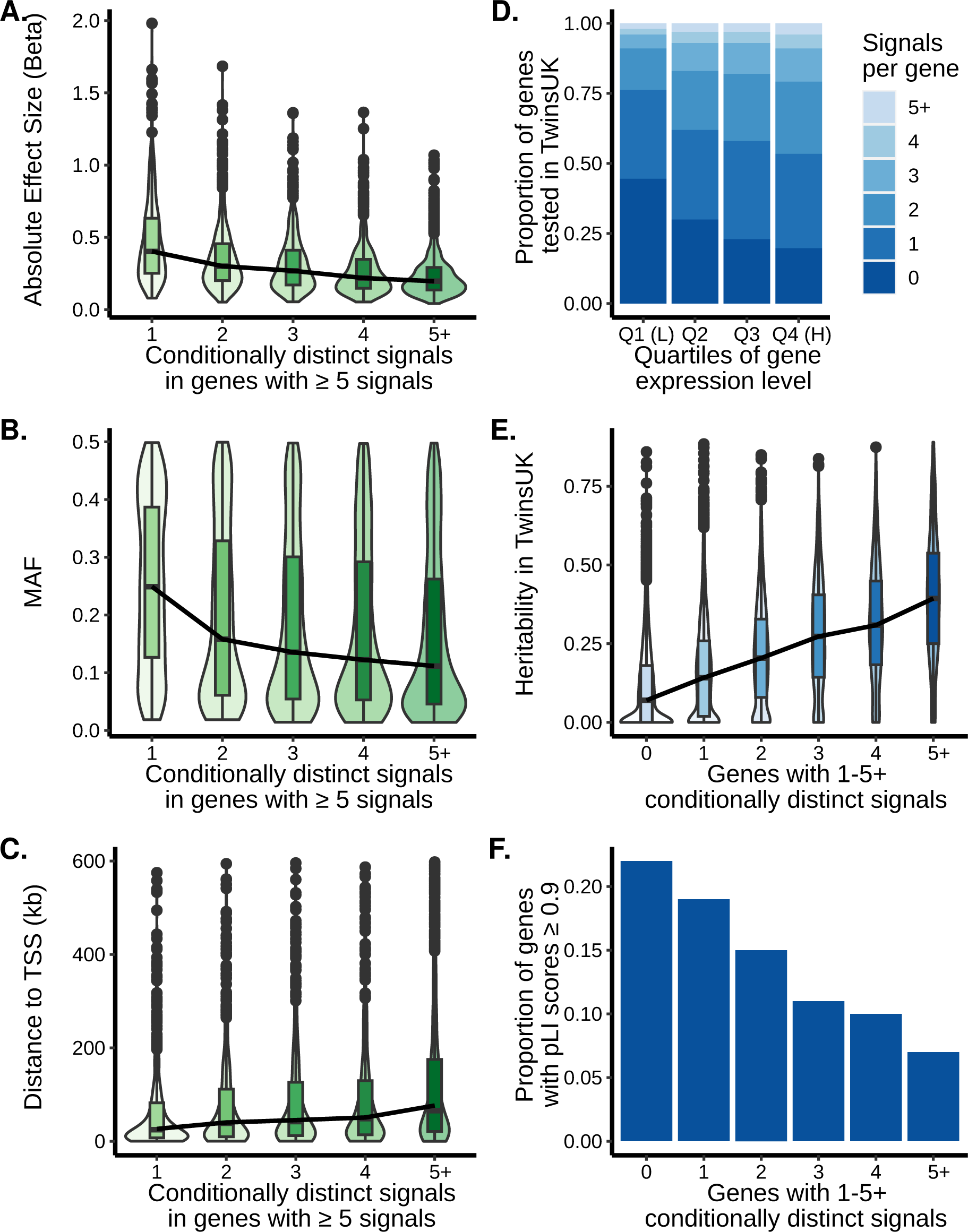
Characteristics of eQTL variants and genes according to the number of significant eQTL signals. Violin plots with inset boxplots of the (A) absolute value of the effect sizes of lead variants, (B) MAF, and (C) distance of the lead variants to the gene TSS for the indicated signals in order of discovery. Only the 661 genes with 5 or more signals were included. For the boxplots, the center line represents the median value, the box limits represent the upper and lower quartiles, whiskers represent the 1.5x interquartile range, and the black circles represent outliers. The black lines connect the median values of each signal group. In C, 163 points with a distance to TSS greater than 600 were excluded. See Figure S6 for genes with one to four eQTL signals. (D) Proportion of genes in TwinsUK with the specified number of eQTL signals separated by gene expression quartiles. Quartile 1 indicates the genes with the lowest expression. The darkest blue are the genes without an eQTL signal and the lightest blue are genes with five or more eQTL signals. (E) Violin plots with inset boxplots of the heritability of genes with the specified number of eQTL signals in TwinsUK. For the boxplots, the center line represents the median value, the box limits represent the upper and lower quartiles, whiskers represent the 1.5x interquartile range, and the black circles represent outliers. The black lines connect the median values of each signal group. (F) Proportion of genes for each signal number with a pLI score ≥ 0.9 out of the total number of genes that have pLI scores available for that signal number.

We next assessed eQTL gene expression levels, heritability, and the probability of the gene being intolerant of loss-of-function variants (pLI)^33^. Genes in the lowest quartile of expression levels made up the smallest proportion of multi-signal genes (23%), while genes in the highest quartile of expression levels contributed to the largest proportion of multi-signal genes (46%; *P* = 0.002; **Figure 3D**). We estimated heritability using the twin structure of the TwinsUK study and determined that eQTL genes had higher expression heritability (median heritability estimate 0.19) than non-eQTL genes (median heritability estimate 0.07) (*P* ≤ 2e-16; **Table S11**), and genes with more eQTL signals showed higher heritability (**Figure 3E**). This trend persisted when genes were separated into quartiles of expression levels, suggesting that genes with higher heritability have more identified eQTL signals independent of the expression level of the gene (**Figure S7**). Lastly, we estimated how tolerant the eQTL genes were to protein-truncating variation based on their pLI scores from GnomAD^33^. Of 12,643 eQTL genes with available pLI scores, 10,625 (84%) were tolerant of truncating variants (pLI < 0.9). eQTL genes with few eQTL signals were more likely to be intolerant of truncating variants than genes with more eQTL signals (**Figure 3F**). For each expression level quartile, the proportion of genes with multiple signals was substantially lower for genes with pLI ≥0.9 than for genes with pLI <0.9. This trend was particularly pronounced in the highest expression category which also has the highest proportion of genes with pLI ≥0.9 (**Figure S8**). We observed the same gene expression and pLI score trends using METSIM gene expression level quartiles (**Figure S9**). Thus, we identified more eQTL signals in highly expressed, more heritable genes that were more tolerant to loss-of-function variants. Higher expression level may be a proxy for power to detect eQTL signals, while higher heritability may reflect a more limited contribution of the environment or technical variation in expression quantification.

### Adipose eQTL identify genes for cardiometabolic trait GWAS signals

To predict candidate genes for GWAS signals, we performed colocalization of conditionally distinct adipose eQTL signals with conditionally distinct GWAS signals for 28 cardiometabolic traits^34–42^ (see Methods)(**Table S12**). We identified 3,605 eQTL signals for 1,861 unique genes that colocalized with signals from at least one GWAS trait (**Table 2; Table S13-15**). All colocalized GWAS-eQTL signals can be visualized using our interactive colocalization browser: https://adipose.colocus.app/. The ten traits with the largest number of eQTL-GWAS signal colocalizations were high-density-lipoprotein cholesterol (HDL-C), log-transformed triglycerides (logTG), total cholesterol (TC), body mass index (BMI), low density lipoprotein cholesterol (LDL-C), waist-to-hip ratio adjusted for BMI (WHRadjBMI), non-HDL-C cholesterol (nonHDL-C), hip circumference (HC), waist-to-hip ratio (WHR), and diastolic blood pressure (DBP) (**Table 2; Table S14**). Among the colocalized eQTL and GWAS signals, only 31% correspond to the gene nearest to the GWAS signal (**Table S14**). On average, 34% of GWAS signals for these 28 cardiometabolic traits had at least one colocalized eQTL signal (**Table S15**). For traits expected to be more relevant to adipose tissue, such as the ratio of abdominal subcutaneous and gluteofemoral adipose tissue volume, 63% of GWAS signals (10 of 16) colocalized with an adipose eQTL signal (**Table S15**). The number of cardiometabolic trait signals with a colocalized eQTL in this meta-analysis is four times greater than the number of results from similar analyses in the METSIM (N) study alone when using the same LD threshold (r^2^ ≥ 0.8)^6^. Thus, larger eQTL studies can identify colocalized eQTL genes for more GWAS signals.

**Table 2:** Summary of GWAS signals colocalized with adipose eQTL signals.

| GWAS trait | GWAS signals | eQTL genes | Total colocalized signal pairs | Primary eQTL signals | Non-primary eQTL signals (percentage of colocalized) |  |
| --- | --- | --- | --- | --- | --- | --- |
| High density lipoprotein-C | 213 | 344 | 369 | 237 | 132 | 36% |
| Triglycerides (log) | 213 | 372 | 393 | 264 | 129 | 33% |
| Total cholesterol | 181 | 294 | 297 | 204 | 93 | 31% |
| Body mass index | 164 | 266 | 272 | 186 | 86 | 32% |
| Low density lipoprotein-C | 154 | 231 | 236 | 153 | 83 | 35% |
| Waist-to-hip ratio adjBMI | 146 | 225 | 238 | 161 | 77 | 32% |
| Non- High density lipoprotein | 142 | 251 | 263 | 188 | 75 | 29% |
| Hip circumference | 124 | 200 | 207 | 150 | 57 | 28% |
| Waist-to-hip ratio | 105 | 176 | 186 | 125 | 61 | 33% |
| Diastolic blood pressure | 100 | 152 | 158 | 116 | 42 | 27% |
| Waist circumference | 87 | 170 | 174 | 116 | 58 | 33% |
| Pulse pressure | 81 | 119 | 120 | 88 | 32 | 27% |
| Type 2 diabetes | 81 | 152 | 160 | 115 | 45 | 28% |
| Systolic blood pressure | 73 | 131 | 135 | 91 | 44 | 33% |
| Coronary artery disease | 69 | 104 | 105 | 71 | 34 | 32% |
| Hemoglobin A1c | 23 | 40 | 40 | 29 | 11 | 28% |
| Gluteofemoral AT adjBMI | 19 | 33 | 34 | 24 | 10 | 29% |
| Fasting glucose | 17 | 29 | 29 | 19 | 10 | 34% |
| Fasting insulin | 16 | 28 | 28 | 17 | 11 | 39% |
| Visceral/Subcutaneous | 14 | 34 | 35 | 23 | 12 | 34% |
| Visceral AT adjBMI | 12 | 29 | 30 | 20 | 10 | 33% |
| Visceral/Gluteofemoral | 12 | 25 | 25 | 20 | 5 | 20% |
| Subcutaneous AT adjBMI | 10 | 22 | 22 | 16 | 6 | 27% |
| Subcutaneous/Gluteofemoral | 10 | 25 | 25 | 19 | 6 | 24% |
| Gluteofemoral AT | 8 | 14 | 15 | 10 | 5 | 33% |
| 2-hour glucose | 3 | 3 | 3 | 2 | 1 | 33% |
| Visceral AT | 2 | 4 | 5 | 3 | 2 | 40% |
| Subcutaneous AT | 1 | 1 | 1 | 1 | 0 | 0% |
| **Total** | **2,080** | **3,474** | **3,605** | **2,468** | **1,137** | **32%** |
| **Total unique** | **--** | **1,861** | **--** | **--** | **--** | **--** |

GWAS signals colocalized with eQTL signals based on lead variant LD $r^2 \geq 0.5$ and coloc PP4 $\geq 0.5$ . GWAS signals indicates the number of unique GWAS signals that are colocalized with at least one eQTL signal. eQTL genes indicates the unique eQTL genes colocalized with at least one GWAS signal. The total colocalized signals indicates the sum of primary and non-primary GWAS-eQTL signal pairs. The total unique row is the total number of unique eQTL genes and signals colocalized with a GWAS signal. C, cholesterol; AT, adipose tissue.

We assessed the colocalized conditionally distinct GWAS-eQTL signals for evidence of gene expression mediating the effect of a genetic variant on a trait using summary Mendelian randomization (SMR)^10^; 2,860 of the 3,587 (80%) analyzed signals had evidence of mediation (*P* < 1.4e-5) (**Table S16**). The subset of signals with evidence of mediation may be more likely to act via those genes to influence the traits.

We next evaluated the contribution of primary versus non-primary signals to GWAS colocalization. We observed 2,468 primary eQTL signals for 1,373 genes and 1,137 non-primary eQTL signals for 596 genes that colocalized with at least one GWAS signal. Inclusion of the non-primary eQTL signals increased the number of GWAS-colocalized signals by 46%. The proportion of eQTL signals that colocalized with at least one GWAS signal was highest for primary eQTL signals and lower for each successively detected eQTL signal, even when accounting for eQTL signal strength (**Figure S10; Table S17**). However, colocalizations for 488 of these 596 genes were only detected using non-primary signals (**Table 2; Table S14**). Overall, the analysis of non-primary eQTL greatly increased the number of GWAS colocalizations.

Many previous studies have performed colocalization with un-conditioned, ‘marginal’ eQTL and GWAS summary statistics. To directly compare the differences between using marginal and conditional results, we also performed colocalization using the marginal eQTL and GWAS statistics. Colocalization analyses with marginal GWAS and eQTL signals identified 1,073 colocalized genes (**Table S18**), 89 of which were detected only in the marginal analysis. Colocalization analyses of the conditionally distinct signals identified 864 (47%) additional genes, 666 of which have multiple eQTL signals (**Table S18**). These results are consistent with previously described limitations of colocalization analysis when marginal eQTL results are used at loci with multiple signals^7,43,44^. These results demonstrate the importance of using conditionally distinct signals to identify GWAS candidate genes, yet suggest that analyses of marginal, unconditioned loci may still provide some value at complex multi-signal loci.

We also colocalized male and female eQTL signals with male and female GWAS signals for a set of sex-biased cardiometabolic traits^37,38,42^, including WHRadjBMI, WC, HC, and body fat distribution^37,38,42^. We identified 144 GWAS-eQTL colocalizations in females and 71 in males (**Table S19-S20**). Of the 138 GWAS-eQTL colocalized signals for WHRadjBMI in only one sex, 82 do not have a corresponding GWAS-eQTL colocalization in the sex-combined analysis. For example, a female eQTL signal at *ADORA1* colocalized with WHRadjBMI in females (**Figure 4**). *ADORA1* encodes an adenosine receptor that suppresses lipolysis in adipocytes, and loss of the receptor leads to glucose intolerance in obese mice^45^. Although the sex-stratified eQTL analysis has limited power, we were able to identify 144 candidate genes for male and/or female GWAS signals, one-third of which were not found in the corresponding sex-combined studies.

**Figure 4.**
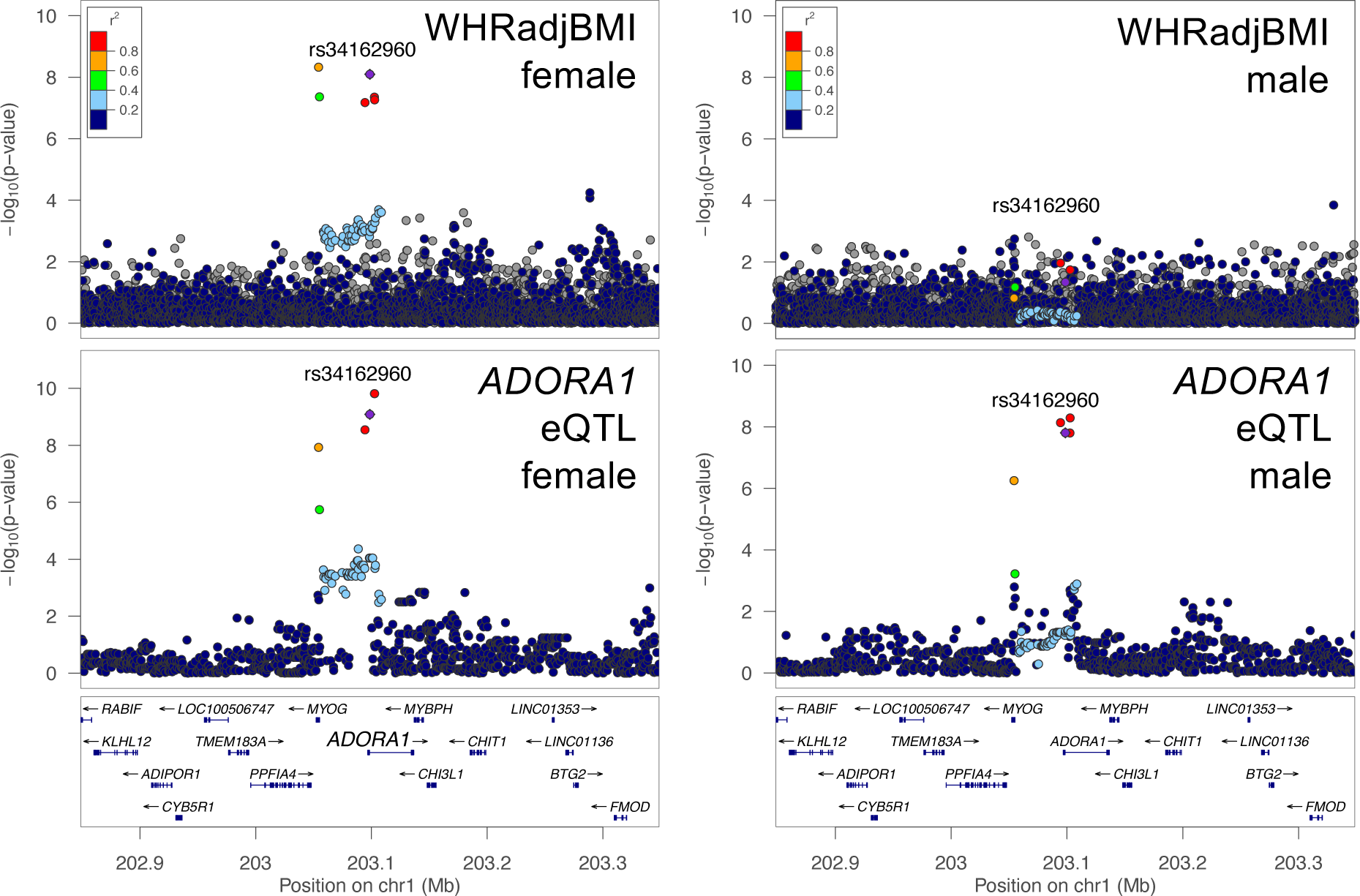
Sex-stratified WHRadjBMI GWAS and *ADORA1* eQTL signal plots. (A) LocusZoom plots for WHRadjBMI female GWAS signal and (B) *ADORA1* female eQTL signal. (C) LocusZoom plots for WHRadjBMI male GWAS signal and (D) *ADORA1* male eQTL signal. All plots are colored by LD with the female GWAS lead variant represented by a purple diamond.

Multiple eQTL signals for the same gene, termed allelic series, that colocalize with multiple GWAS signals from the same trait can provide additional confidence that the gene influences the trait. In the eQTL meta-analysis, 33 unique genes harbored allelic series that colocalized with allelic series for at least one GWAS trait, corresponding to 144 of 3,605 (4%) GWAS-eQTL colocalized signal pairs (**Table S14; Table S21**). We used only the 30 genes that harbored nearly independent eQTL signals (LD r^2^ < 0.05) to estimate causal effects using MRLocus;^46^ all eQTL signals for the gene, including those that did not colocalize with GWAS signals, were included in the MR analysis. Among the 30 genes, 21 have evidence of mediation (adjusted *P* ≤ 0.25) (**Table S21; Figure S11**). For example, *ZNRF3* has two eQTL signals that are colocalized with two WHRadjBMI GWAS signals (**Figure 5A; Figure S12**). The alleles associated with lower WHRadjBMI at both signals were associated with higher *ZNRF3* expression levels, as displayed by a negative GWAS vs eQTL slope from MRLocus (adjusted *P* = 0.18; **Figure 5B; Table S21**). The two signals provide evidence for an estimated gene-to-trait effect of -0.19, indicating that increasing adipose *ZNRF3* expression level by one population standard deviation should reduce WHRadjBMI by 19% of its population standard deviation. For further support, the observed trait-gene association in METSIM shows higher *ZNRF3* expression level associated with lower WHR (*P* = 0.04; beta = -0.85; **Figure 5C**), although this association may be confounded by factors that influence both *ZNRF3* and WHR, or reverse causal effects. *ZNRF3* encodes a membrane-bound E3 ubiquitin ligase, which is a receptor for R-spondins and functions as a negative feedback regulator in the WNT signaling pathway. ^47,48^ When we further limit the allelic series to pairs of signals for which LD D’ < 0.1, 9 genes had independent allelic series and 7 of them showed evidence of mediation (**Table S21**). For example, *PDE3A* has four eQTL signals that colocalized with four HDL-C GWAS signals (**Figure 5D-F; Figure S13**). For all four signals, the alleles associated with lower HDL-C were associated with higher *PDE3A* expression level. Two of the signals are nearly independent (lead variants pairwise LD r^2^<0.05, D’ <0.1) and provide evidence for an estimated gene-to-trait effect of -0.14 (adjusted *P* = 0.15; **Figure 5E**). PDE3A regulates cAMP signaling and has been shown to have higher expression in the hearts of diabetic than non-diabetic rats.^49,50^ Colocalized allelic series of GWAS and eQTL signals provide stronger confidence that gene expression in the assayed tissue influences the trait, and gene-based dosage effects may help predict the impact that therapies modulating a gene will have on traits.

**Figure 5.**
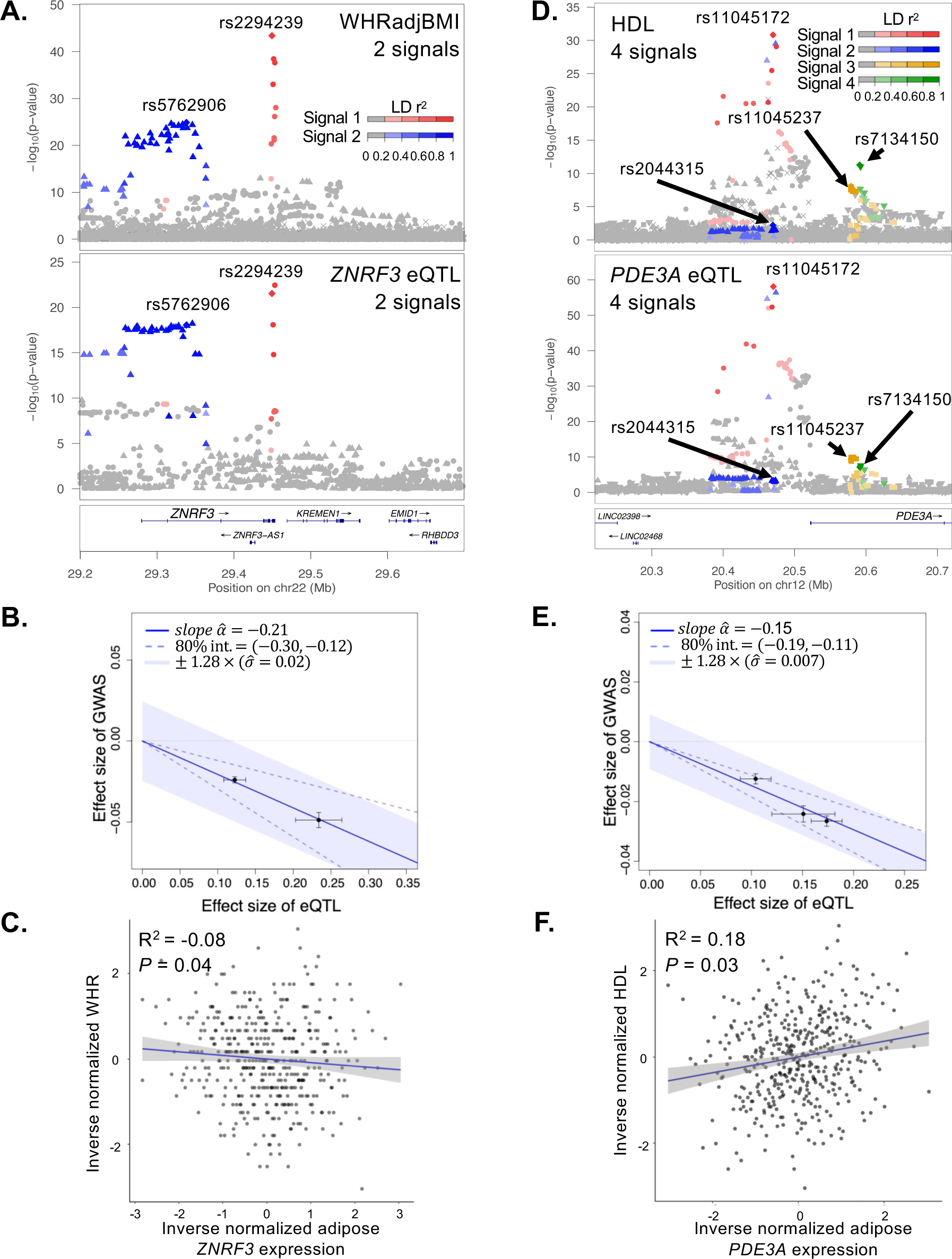
Colocalization of two or more GWAS signals with two or more eQTL signals at *ZNRF3* and *PDE3A*. A. LocusZoom plots of WHRadjBMI GWAS summary statistics (Pulit et al 2019) (top) and marginal *ZNRF3* eQTL data for the meta-analysis (bottom). Both plots show two signals colored by the GWAS lead variants (red diamond, 1^st^ signal chr22:29,449,477, rs2294239; blue diamond, 2^nd^ signal chr22:29,338,235, rs5762906). The red circles and blue triangles indicate genetic variants in stronger LD with the 1^st^ or 2^nd^ signal, respectively and are shaded based on LD. Signal 1 in the GWAS is colocalized with signal 1 of the eQTL dataset (LD r^2^ = 0.90; coloc PP4 = 0.99) and signal 2 for both datasets are also colocalized (LD r^2^ = 1.00; coloc PP4 = 0.98). B. Effect sizes of the WHRadjBMI GWAS signals (y-axis) versus the effect sizes of the *ZNRF3* eQTL signals (x-axis) from MRLocus. Each point represents a colocalized eQTL signal with standard error bars. The solid blue line represents the slope of the effect of the gene on the trait, and dotted blue lines represent the confidence interval. The slope estimates a gene-to-trait effect of -0.19, meaning that increasing adipose *ZNRF3* expression level by one population standard deviation should reduce WHRadjBMI by 19% of its population standard deviation. C. Scatter plot of inverse normalized waist-to-hip ratio (x-axis) and *ZNRF3* gene expression (y-axis) in METSIM (S) (n = 420). Each point represents an individual sample, the blue line represents the linear regression slope and the 95% confidence interval is shown in gray. The correlation value and association *P*-value are shown. D. LocusZoom plot of the HDL-C GWAS summary statistics (Graham et al 2021) (top) and marginal *PDE3A* eQTL data for the meta-analysis at (bottom). Both plots show four signals colored by the GWAS lead variants (red diamond, 1^st^ signal chr12:20,470,221, rs11045172; blue diamond, 2^nd^ signal chr12:20,470,009, rs2044315; yellow diamond, 3^rd^ signal chr12:20,579,083, rs11045237; green diamond, 4^th^ signal chr12:20,591,332, rs7134150). The red circles, blue triangles, yellow squares, and green inverted triangles indicate genetic variants in stronger LD with the 1^st^, 2^nd^, 3^rd^, or 4^th^ signal, respectively and are shaded based on LD. Signal 1 in the GWAS is colocalized with signal 1 of the eQTL dataset (LD r^2^ = 1.00; coloc PP4 = 1.00), signal 2 for the GWAS is colocalized with signal 4 of the eQTL dataset (LD r^2^ = 0.93; coloc PP4 = 1.00), signal 3 for the GWAS and signal 2 for the eQTL dataset are colocalized (LD r^2^ = 0.42; coloc PP4 = 1.00), and signal 4 for the GWAS and signal 3 for the eQTL dataset are colocalized (LD r^2^ = 0.94; coloc PP4 = 0.99). E. Effect sizes of the HDL-C GWAS signals (y-axis) versus the effect sizes of the *PDE3A* eQTL signals (x-axis) from MRLocus. Each point represents a colocalized eQTL signal with standard error bars. The solid blue line represents the slope of the effect of the gene on the trait, and dotted blue lines represent the confidence interval. The slope estimates a gene-to-trait effect of -0.14, meaning that increasing adipose *PDE3A* expression level by one population standard deviation should reduce HDL-C by 14% of its population standard deviation. F. Scatter plot of inverse normalized HDL-C (x-axis) and *PDE3A* gene expression (y-axis) in METSIM (S) (n = 420). Each point represents an individual sample, the blue line represents the linear regression slope and the 95% confidence interval is shown in gray. The correlation value and association *P*-value are shown.

### Regulatory variants within eQTL signals

To predict the genomic features that may be responsible for eQTL signals, we investigated the location of eQTL variants relative to adipose chromatin states. We compared enrichment of conditionally distinct eQTL signals (lead and proxy variants r^2^>0.8) relative to signals for genes without an eQTL in Roadmap Epigenomics adipose tissue promoters and enhancers based on the order signals were discovered in the stepwise conditional analysis.^51^ The 1^st^ through 4^th^ eQTL signals were significantly enriched in promoters and enhancers, whereas the 5^th^ and higher eQTL signals were not (**Figure 6A; Figure S14; Table S22**). Primary eQTL signals were much more strongly enriched in promoters (odds ratio = 3.5) than enhancers (odds ratio = 2.2). 2^nd^ through 4^th^ signals were slightly more enriched in promoters than enhancers and each signal showed sequentially decreasing enrichment levels (**Figure 6A; Figure S14; Table S22**). These results show that non-primary signals are less often located in promoters and increase the total number of signals detected in both promoters and enhancers.

**Figure 6.**
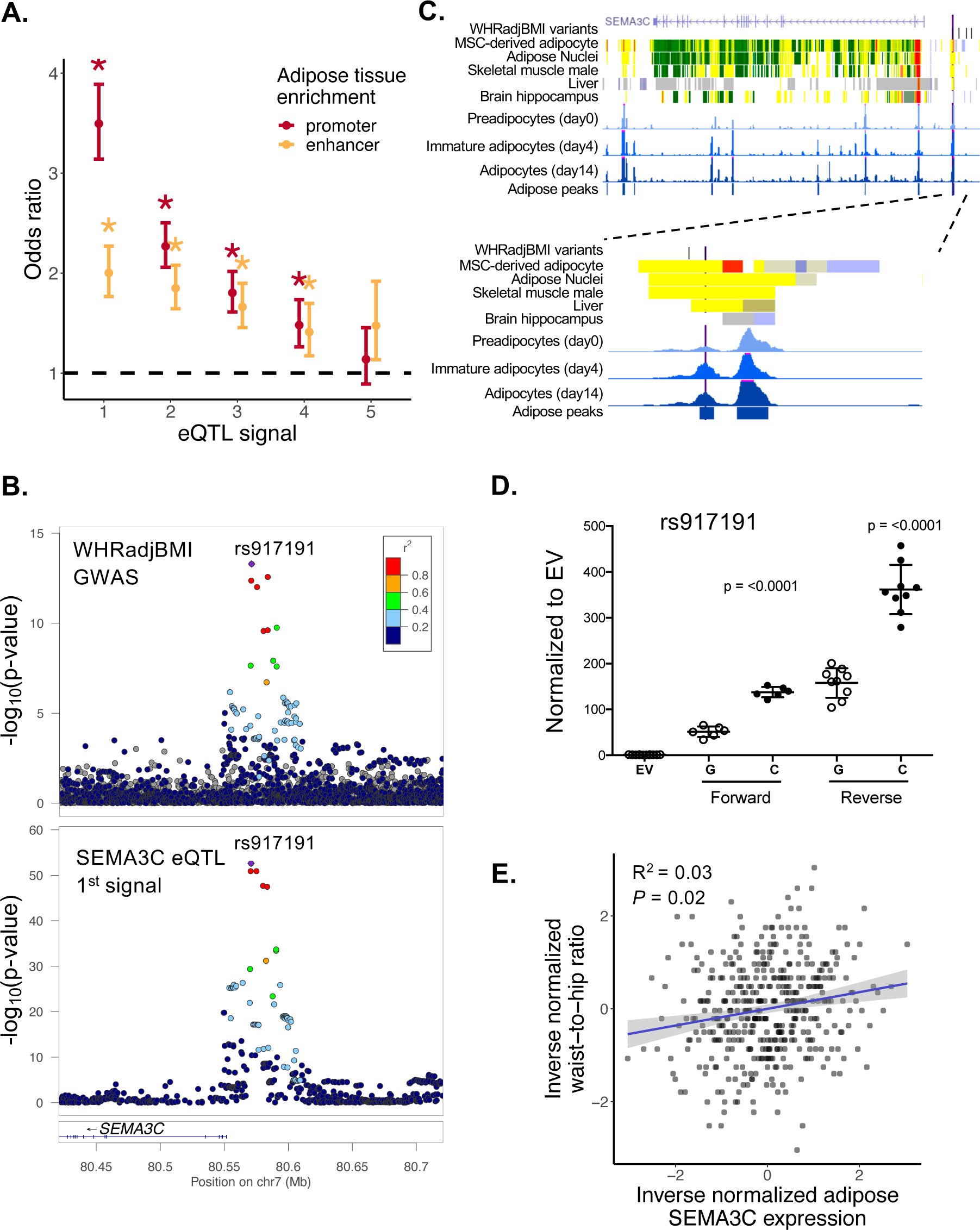
Regulatory annotation enrichment of eQTL signals and validation of allelic effects on transcriptional activity at *SEMA3C*. A. eQTL signals enriched in Roadmap Epigenomics chromatin states in adipose tissue compared to genes without an eQTL separated by signal number. Dark red represents promoters and gold represents enhancers. The bars represent the upper and lower 95% confidence intervals. The asterisk represents significant Bonferroni-adjusted enrichment values that do not overlap an odds ratio (OR) of 1 (black dashed line). B. LocusZoom plots of the WHRadjBMI GWAS summary statistics (Pulit et al 2019)(top) and the *SEMA3C* meta-analysis eQTL data conditioned on all but signal 1 (bottom). Both plots show the same lead variant represented by a purple diamond (chr7:80,570,871; rs917191). Other variants are colored based on the LD r^2^ 1000G EUR with the lead variant. Signal 1 in the GWAS dataset is colocalized with signal 1 of the eQTL dataset (LD r^2^ = 1.0; coloc PP4 = 1.0). C. UCSC genome browser tracks showing regulatory annotations that overlap *SEMA3C* eQTL variants. In the *SEMA3C* SNPs track, the lead variant is shown in purple and proxy variants (LD r^2^ ≥ 0.8) are shown in black. The chromHMM tracks are from Epigenomic Roadmap for mesenchymal stem cell-derived adipocytes, adipose nuclei, skeletal muscle, liver, and brain hippocampus; red represents a promoter-like signature, yellow represents an enhancer-like signature, green represents a signature for elongating RNA polymerase, and gray represents low to no signal. The blue signal tracks represent ATAC-seq accessible chromatin in SGBS cells at differentiation day 0, day 4, and day 14. The METSIM adipose peaks are ATAC-seq peaks detected in at least 3 adipose tissue samples. *SEMA3C* gene annotations are from UCSC genes. The bottom figure shows the browser tracks zoomed in to the region around rs917191. D. Relative transcriptional activity of rs917191-G and rs917191-C in hWAT adipocytes from dual-luciferase reporter assays. Values indicate transcriptional activity relative to an empty vector (EV), points represent independent clones with standard error bars, and *P*-values from Student’s unpaired t-tests compare activity between alleles. E. Scatter plot of inverse normalized waist-to-hip ratio (y-axis) and *SEMA3C* gene expression (x-axis) in METSIM (S) (n = 420). Each point represents an individual sample, the blue line represents the linear regression slope and the 95% confidence interval is shown in gray. The correlation value and association *P*-value are shown.

To identify candidate regulatory variants that may act through adipose regulatory elements, we compared eQTL variants to sites of accessible chromatin defined by ATAC-seq peaks in adipose tissue and preadipocytes and mature adipocytes of the human Simpson Golabi Behmel Syndrome (SGBS) cell strain.^52^ Of the 34,438 eQTL signals, 40% had at least one proxy variant located in an adipose tissue accessible chromatin region, and 51% had at least one variant in a mature adipocyte region (**Table S23**). Among the eQTL signals colocalized with GWAS signals, 60% and 72% had at least one variant in adipose tissue or mature adipocyte accessible chromatin, respectively (**Table S23**). Among ∼16K chromatin regions more accessible in adipocytes than preadipocytes,^53^ adipose eQTL enrichment was significant for the 1^st^ through 3^rd^ signals (odds ratio for primary signals = 1.6) while no signals were significantly enriched in ∼18K chromatin regions more accessible in preadipocytes than adipocytes^53^ (odds ratio for primary signals = 1.0; **Figure S14; Table S22**). Thus, more than half of meta-analysis eQTL signals contain plausible regulatory variants located in regions of adipose or adipocyte accessible chromatin.

We further investigated potential regulatory variants at one colocalized GWAS-eQTL signal. The primary *SEMA3C* eQTL signal colocalized with a WHRadjBMI GWAS signal (LD between lead variants, r^2^ = 1.0; coloc PP4 = 1.0) (**Figure 6B**). *SEMA3C* is an adipokine predominantly expressed in mature adipocytes and regulated by weight changes.^54^ The lead variant (chr7:80,570,871; rs917191) is located in an accessible chromatin region in both adipose and adipocytes^53^ (**Figure 6C**, **Table S24**), while a variant in high LD with the lead variant (chr7:80,580,219; rs12537553, r^2^ = 0.89) is located in an accessible chromatin region in muscle^55^. We tested both variants for allelic differences in transcriptional activity in preadipocytes and differentiated adipocytes from hWAT and SGBS cell lines, as well as myoblasts and differentiated myocytes from the LHCN-M2 cell line. rs917191 showed strong enhancer activity and 2.3-to 6.2-fold higher transcriptional activity for the C allele in preadipocytes, adipocytes, myoblasts, and myocytes, whereas the proxy variant rs12537553 showed no significant differences in activity (**Figure 6; Figure S15-S16**). The rs917191-C allele was associated with increased WHRadjBMI risk, higher *SEMA3C* gene expression levels, and greater transcriptional activity than the rs917191-G allele. The trait-gene association in METSIM also indicates that higher *SEMA3C* expression is associated with higher WHR (*P* = 0.02; beta = 0.11; **Figure 6**). These data suggest that rs917191 may alter *SEMA3C* activity in adipose tissue and lead to effects on WHR. The hundreds of other colocalized GWAS and eQTL signals suggest that many additional regulatory mechanisms responsible for GWAS signals may be discovered (**Table S14**).

## Discussion

We carried out the largest adipose tissue eQTL study to date and present a broadly applicable framework to efficiently define conditionally distinct eQTL signals across multiple studies. We detected 34K conditionally distinct eQTL signals in 18K genes, which is 2.3-fold more signals and >1.6-fold more eQTL genes than detected by any of the five studies alone. On average, each gene had ∼2 eQTL signals, and some highly expressed genes harbored as many as 10 signals. Colocalization of eQTL with GWAS identified 1,861 candidate genes for over 2,000 cardiometabolic trait GWAS signals across 28 traits, at least 4-fold more than any previous adipose eQTL study when accounting for differences in LD threhsolds.^6^ Including non-primary eQTL signals enabled discovery of 46% more GWAS-eQTL colocalized signals than using primary signals alone, suggesting that current, widely used eQTL studies remain underpowered and that non-primary eQTL signals can help explain some of the “missing regulation.”

The newly identified eQTL signals identified more distal variant effects on expression. Previous studies by us and others have shown that non-primary eQTL signal lead variants are located further away from the gene transcription start sites than primary eQTL lead variants.^6,15^ We show that this trend continues with additional eQTL signals and that the median distance from variant to gene TSS between 1^st^ and 5^th^ signals increases 2.8-fold. In addition, eQTL variants for 2^nd^, 3^rd^, and 4^th^ signals showed successively less enrichment in adipose promoters and enhancers, especially for promoters, consistent with other studies^23^ and the hypothesis that primary eQTL tend to act on promoters. The non-primary eQTL signal distances to TSS are thus more like GWAS signals, suggesting that a larger proportion of non-primary eQTL would colocalize with GWAS signals; however, primary eQTL showed more GWAS colocalizations, which may reflect still limited power to detect eQTL.

The conditionally distinct signals also provided a more thorough understanding of gene regulation. Although a prior study showed consistent effect sizes among primary and non-primary eQTL signals,^15^ in our previous study^6^ and here we observed that effect sizes for 1^st^ signals were twice as large as those from 5^th^ signals, which is expected because variants with stronger effects on a trait are easier to detect against a background of other genetic and environmental factors. We also found that the median heritability for genes with five or more signals was 2.5-fold higher than genes with only one signal, consistent with a study of blood eQTL.^12^

We found that genes with high levels of intolerance of loss-of-function mutations are less likely to have multiple signals than those with lower levels of constraint, as shown previously in brain tissue^15^. For genes in the highest quartile of expression we observed two opposing forces that affected the probability of detecting an eQTL. Genes in the higher quantiles of expression have greater power to be detected as eQTLs due to higher read counts, however genes in the higher quantiles of expression are also substantially more likely to have low tolerance of loss-of-function mutations, thus decreasing the power to detect eQTLs. Overall, using a larger, better-powered eQTL study allowed us to more comprehensively dissect gene regulation.

Integration of GWAS, eQTL, and regulatory elements helped identify plausible regulatory mechanisms. Over 1,800 eQTL genes colocalized with GWAS signals, and 72% of the colocalized signals had lead or proxy variants (LD r^2^≥0.8) located in mature adipocyte accessible chromatin regions, providing candidate regulatory variants, including a variant we validated by showing allelic differences in transcriptional activity at *SEMA3C*. One challenge of discovering more eQTL and colocalizations is that cardiometabolic GWAS signals can show evidence of colocalization with eQTL for more than one gene, even if genetic effects on each gene do not affect the GWAS trait. To address this challenge, we examined mediation using MRLocus on the subset of genes for which two or more apparently independent eQTL signals (LD r^2^< 0.05) colocalized with two or more GWAS signals. This analysis provided stronger evidence of causal effects for 21 genes and estimates of their gene-based dosage effects on the GWAS trait. Despite our desire to analyze pairs of independent colocalized signals, 80% of the 70 signal pairs tested for mediation still have D’> 0.1, suggesting that haplotype effects may still influence gene dosage estimates. Nonetheless, evidence of mediation and estimates of the dosage effect of genes on traits strengthens the support for targeting a gene with drug therapeutics to ameliorate disease.

Although this study of >2,000 individuals is relatively large, it still has limitations. Continued increase in sample size should identify additional signals, even after the number of detected eQTL genes reaches saturation. Expanded studies in more diverse populations would enable analysis of additional variants and thus detection of additional eQTL signals. In addition, our sex-stratified eQTL meta-analyses were underpowered (270 female and 418 male individuals), and additional sex-dependent eQTL remain to be identified. Lastly, we identified eQTL in bulk adipose tissue, which integrates the eQTL signals across cell types; we may not have detected some cell-type-specific eQTL. Future eQTL discovery from single cells or nuclei are needed to distinguish these cell type effects.

In summary, this adipose eQTL analysis tripled the size of previous studies, furthered understanding of allelic heterogeneity in gene regulation, greatly expanded discovery of eQTL colocalized with cardiometabolic trait GWAS signals, and identified thousands of candidate genes that may lead to new drug therapies.

## Online Methods

### Study cohorts, quality control, and RNA-sequencing

#### METSIM

The sample collection and genotyping of 10,197 male individuals from Kuopio, Finland in the METabolic Syndrome In Men (METSIM) study was described previously.^28,56^ Subcutaneous adipose tissue was sampled from near the umbilicus for two non-overlapping sets of samples for which the RNA-sequencing was performed at separate times. One subgroup, referred to as METSIM (N), has 426 participants who provided a needle tissue biopsy for which RNA-seq was previously described.^6^ Compared to the previous report, we removed eight samples from individuals who also participated in the FUSION study described below. The second subgroup, referred to as METSIM (S), has 420 participants who provided surgical biopsies; these individuals are independent from METSIM (N) and FUSION, and the RNA-seq was described previously, although this is the first report of eQTL.^57^ Briefly, for both METSIM (N) and METSIM (S), we removed adaptor sequences and sequences with phred quality scores of < 20 using Fastx-toolkit^58^ and Cutadapt^59^ (v.1.18) respectively, as described.^6^ We aligned RNA-seq reads to the hg19 reference genome using STAR^60^ for both METSIM (N) (v. 2.4.2a) and METSIM (S) (v. 2.7.3a) as described.^6^

#### FUSION

Inclusion criteria for the Finland-United States Investigation of NIDDM (FUSION) tissue collection has been described previously.^30,55^ Genotyping of FUSION tissue biopsy participants has been described previously.^55^ Briefly, we collected subcutaneous adipose tissue samples from near the umbilicus using surgical biopsy.^30^ We followed the same procedures of RNA extraction and mRNA-seq, and quality control (QC) as for muscle.^55^ Subcutaneous adipose tissue sample RIN ranged from 5.1 to 8.8 (median 7.4). RNA-seq reads from 280 subcutaneous adipose tissue samples were aligned to the hg19 reference genome using STAR^60^ (v.2.7.3a).

#### TwinsUK

Sample collection, SNP genotyping, and quality control were conducted as previously described.^30,61^ Genotype data were available for 722 participants for whom adipose tissue gene expression data were available. Subcutaneous adipose tissue RNA was extracted from punch biopsies from a sun-protected area of the abdomen, and RNA sequencing and data processing carried out as described elsewhere.^30,62,63^ RNA-Seq reads were aligned to the hg19 reference genome using STAR^60^ version 2.4.0.1.

#### GTEx

Genotype-Tissue Expression (GTEx) V8 sample collection, whole genome sequencing, RNA-sequencing and quality control for all samples, including subcutaneous adipose tissue, has been described previously.^1^ We obtained dbGaP permissions and accessed the genotype files (phs000424). We subset the subcutaneous adipose tissue samples that had genotype information. In our analysis, we tested two GTEx studies, one with all individuals and the other with only individuals with European ancestry as estimated through use of principal component analysis described in the original study.^1^ We downloaded the previously described reads per gene from the GTEx portal.^1^ We lifted over the variants from hg38 to hg19 using the variant look-up file provided by GTEx and kept the gene assignments as reported.

### Genotype Imputation of array-genotyped samples and inclusion of WGS samples

In studies except the whole genome sequenced GTEx study, samples were imputed using the Haplotype Reference Consortium panel (hg19)^64^ as previously described.^28,30,55,61^ In each study with imputed genotype data, we excluded variants with low imputation quality (R^2^ < 0.3 or 0.5)(**Table S1**). In all studies we excluded variants with MAF <0.01. We coded the X chromosome genotypes as diploid (0/2) for males. For analysis, we retained 6,995,803 variants that were present in all five studies.

### Gene level quantification and ADIPOQ expression-based sample inclusion

For all studies except GTEx, to quantify the read counts per gene, we used GENCODE v19^65^ as the reference and the quan function from QTLtools package^19^ (METSIM and TwinsUK) or QoRTs^66^ (v.1.3.6) (FUSION). In each study, to select for more highly expressed genes, we retained genes with 5 or more counts in at least 25% of the individuals in each study. We calculated counts per million (CPMs) normalized for library size by adjusting the CPMs by the Trimmed mean of M-values (TMM)^67^ using edgeR^68^ (v.3.36.0). We included subcutaneous adipose tissue samples that had >150 CPM for *ADIPOQ* gene expression (**Figure S2**), an arbitrary threshold we used as a proxy for substantial adipocyte content. The total sample sizes used for analysis are given in **Tables 1, S1, and S2.**

### Study-level eQTL analysis

In each study, we inverse normalized the gene expression values. To account for technical and physiological differences across samples we constructed probabilistic estimation of expression residuals (PEER) factors^69^ using the inverse-normalized gene expression. For all studies except TwinsUK (see below), we performed linear regression of expression values with BMI as a covariate to remove the effect of BMI from the residuals (which will remove the effect of BMI from those captured by the subsequent PEER factor analysis). To account for unknown technical variation we generated PEER^69^ using the gene expression residuals adjusted for BMI. In each study, to identify the number of PEER factors to include as covariates in our eQTL analysis, we generated PEER factors in sets of 10 from 0 to 100 and performed eQTL analysis for each PEER factor set using the ordinary least squares local-eQTL analysis from APEX.^21^ We calculated the number of significant genes as genes with ≥1 variants with FDR <1%. We quantitated the percent change in the number of significant genes for each successive increase in PEER factors and selected the PEER factor number after which the increase in significant genes was < 1%. For TwinsUK, we used a linear mixed effects regression model of gene expression^70^, including BMI, family zygosity, and SNP genotyping chip to create adjusted gene expression residuals. We then used the same PEER factor generation on the adjusted residuals and selection process described above.

For each study, we performed local-eQTL analysis of variants within 1 Mb of the canonical gene transcription start site using the ordinary least squares analysis in APEX.^21^ The APEX regression model is equivalent to the FastQTL model.^71^ We performed eQTL analysis using inverse normal transformed gene expression, including BMI, study-specific factors, and PEER factors as covariates. We output a summary statistics file for use in subsequent analyses.^21^

### eQTL meta-analysis and conditional meta-analysis

For meta-analysis and in the stepwise conditional meta-analysis, we included variants present in all studies and genes that were expressed in ≥2 studies. We performed inverse-variance weighted meta-analyses in APEX using the study-specific eQTL summary statistics, including either the GTEx-all populations or GTEx-European Americans samples. To enable summary statistic-based conditional analysis, for each study we created a covariate-adjusted genotype variance-covariance matrix (APEX).^21^ Using study-specific summary statistics and covariate-adjusted variance-covariance matrices, we performed sequential rounds of stepwise study-specific conditional analysis followed by inverse-variance weighted meta-analysis to detect a new lead conditionally distinct signal across studies.^21^ Specifically, for genes with at least one variant with a *P* ≤ 1e-6, we performed a forward and backward selection with a threshold of *P* ≤ 1e-6 on the conditional *P*-values. For entry into the model we required lead variants of conditionally distinct signals to have LD r^2^ ≤ 0.7 with the lead variants of the prior signal(s). For eQTL genes with more than one conditionally distinct signal, we extracted the lead variants for the conditionally distinct signals, and for each lead variant, performed eQTL analysis conditioning on all other lead variants (termed all-but-one analysis) as implemented the Apex2R software.^72^ To create comprehensive summary statistics for all eQTL genes (*P* ≤ 1e-6), we combined these all-but-one results for multi-signal genes with the marginal meta-analysis summary results for single-signal genes.

We also performed conditional analysis for each individual study, using the same forward-backward selection procedure, variant inclusion r^2^ criteria, and *P*-value threshold for signal inclusion as for the conditional meta-analysis.

To compare the conditionally distinct lead variants from the meta-analysis including GTEx-European Americans to conditionally distinct lead variants from the meta-analysis with the GTEx-all samples, we estimated the LD r^2^ between all combinations of lead variants for conditionally distinct eQTL signals of the same eQTL gene using PLINK^73^ (v.1.90b3). We used 40,000 unrelated United Kingdom Biobank (UKBB) subjects as the LD reference panel^74^. We considered any lead gene-variant pairs with the same variant or that had an LD r^2^ ≥ 0.8 to be the same signal. All LD look-ups used this UKBB reference panel unless stated otherwise.

### Sex-stratified eQTL meta-analysis and conditional meta-analysis

We performed sex-stratified eQTL meta-analyses in the studies that contained both males and females (FUSION and GTEx-European American). For each study and sex, we identified sets of PEER factors and ran the local-eQTL analysis as described above. We performed sex-specific meta-analysis and conditional meta-analysis as described above.

To ask if we identified the same eQTL signals in females and males, for each gene we estimated the LD r^2^ for all pairs of female and male conditionally distinct lead variants, using PLINK^73^ (v.1.90b3) and UKBB as the LD reference panel. To compare the male and female eQTL effect sizes for each gene, we extracted the female marginal meta-analysis lead variant and the same variant from the male marginal meta-analysis. We repeated the analysis extracting the male marginal meta-analysis lead variant and the same variant from the female marginal meta-analysis. We plotted the effect size comparisons using ggplot2 (v.3.4.0).^75^

### Adipose and blood eQTL comparison

We downloaded the full blood local-eQTL summary statistics from the eQTLGen^32^ website. If the gene had ≥ 1 variant with an eQTL (FDR ≤0.05; *P* < 2e-5), we extracted the lead variant per gene in the blood study. For genes with >1 variant with *P* = 3e-310, we extracted all variants with that *P*-value. For each gene tested in common between the blood and adipose eQTL studies, and for each of the adipose conditionally distinct signals in the gene, we determined the LD r^2^ between each lead conditionally distinct adipose variant and the gene’s lead blood eQTLgen variant using UKBB as a LD reference in PLINK (v.1.90b3).^73^ If eQTLGen contained multiple potential lead variants with a *P* ≤ 3e-310, we chose the variant with the highest LD r^2^ with the adipose eQTL lead variant. We defined shared adipose and blood eQTL signals as those with r^2^ ≥ 0.2 and repeated the analysis with other more stringent LD r^2^ thresholds (≥0.4, 0.6, and 0.8). We considered the blood eQTL signals not shared if the pairwise LD r^2^ < 0.2 between adipose and blood lead variants per gene. We repeated this process using only adipose primary signals or adipose non-primary signals.

### eQTL signal characterization

We compared various characteristics across conditionally distinct eQTL signals and genes. We extracted the conditionally distinct eQTL betas and took the absolute value. We calculated the effect allele frequency (EAF) in METSIM (N), METSIM (S), FUSION, and TwinsUK based on the individuals present in the meta-analysis and in GTEx based on all individuals with genotypes. To estimate the across-study allele frequency, we estimated a sample size-weighted effect allele frequency using METAL^76^ and then used the resulting EAF to calculate the MAF. For each gene, we extracted the initial TSS position from the GENCODE v19^65^ gtf file (gene start for positive strand, gene end for reverse strand). We calculated the distance from the lead conditionally distinct variant to gene TSS by taking the absolute value of the difference between the TSS position and the variant position. To test for differences in eQTL betas, MAF and distance to TSS between pairs of eQTL signal numbers, we used Mood’s median test^77^ which tests for a difference in medians using median_test() in R (v.4.1.3).

We used the LocusZoom software (v.1.4) with the November 2014 1000G EUR reference panel to create all locus plots.^78^ We used marginal eQTL meta-analysis summary results for all plots unless specified as an all-but-one (AB1) plot. The AB1 plots are conditioned on all the signals except the one specified. All other plots were created using ggplot2^75^ (v.3.4.0) in R (v. 4.1.3).

### Heritability estimations

We estimated the heritability of gene expression levels in TwinsUK using 186 dizygotic and 131 monozygotic twin pairs. We calculated the residuals of gene expression level adjusted for technical covariates, including GC mean, median insert size, primer index, date of sequencing and RNA extraction batch. Then, we used the residuals to calculate heritability with twinlm() function from mets package in R (v.4.0.3). We reported the heritability estimated by the ACE model, which assumes the variance in gene expression level to be partitioned into variances of additive genetic factors (A), shared environmental factors (C) between co-twins, and unique environmental factors (E) that are not related between co-twins. Inclusion or exclusion of age as a covariate did not significantly change the results. We tested the relationship between heritability and the number of eQTL meta-analysis signals using linear regression models. As genes with higher expression levels have more statistical power to detect an eQTL or have higher heritability estimates, we also adjusted for the quartile groups of gene expression levels in the model. We calculated the quartiles with the median expression levels among all the samples for each gene within TwinsUK and METSIM (S). We tested for statistical difference between proportions using prop.test() in R.

### pLI scores

We downloaded the gnomAD^33^ (v.2.1.1) loss-of-function metrics by gene table from the gnomAD website. We matched the adipose eQTL genes with the gnomAD table to extract the values for the probability of loss-of-function intolerance (pLI). Then, we calculated the proportion of genes tested for an eQTL with a pLI score ≥ 0.9 out of all genes (with pLI scores available) separated by the number of eQTL signals per gene. We repeated this using only genes expressed in TwinsUK or genes expressed in METSIM (S) and separated the genes by meta-analysis eQTL signal number and TwinsUK or METSIM (S) gene expression quartile (described above).

### GWAS signal identification and conditional analysis

We downloaded GWAS summary statistics for 28 traits from the locations listed in **Table S12**, including sex-stratified GWAS summary statistics for WHRadjBMI^37^, WC^38^, HC^38^, and the fat depot traits.^42^ For each trait, we used GWAS summary data from analysis of European-ancestry individuals.

To meet the coloc assumption of no more than one signal in a region per dataset^43^, we isolated conditionally distinct GWAS signals. When conditionally distinct signals were described, we used the reported lead variants. To isolate each signal, we conditioned on any reported lead variant within 500 kb of the signal of interest. We ran GCTA cojo-cond with default parameters and used the UKBB LD reference panel. This process generated approximate conditional summary statistics for each signal of interest (termed ‘all-but-one’ summary statistics), including variants ±500 kb from the lead variant. If no other lead variant was located within 500 kb of the lead variant of interest, we used the marginal summary statistics as the all-but-one summary statistics.

When conditionally distinct GWAS signals were not reported for a dataset (UKBB traits WC and HC, and GLGC lipid traits), we identified conditionally distinct lead variants from the marginal GWAS summary statistics. We defined a locus as a lead variant (marginal *P* ≤ 5e-8) and its 500 kb flanking regions using swiss (v.1.1.1). If another lead variant was within 1 Mb, we clumped their two loci together into a super-locus. We repeated this clumping until there was greater than 1 Mb between any lead variant in the super-locus and any lead variant in another locus or super-locus. We then used GCTA^20^ cojo-slct (MAF ≥ 1%; collinearity < 0.5; conditional *P* ≤ 5e-8; UKBB LD reference panel) to identify conditionally distinct lead variants within each locus or super-locus. For loci/super-loci with more than one signal, we isolated each signal using GCTA cojo-cond (MAF ≥ 1%; collinearity < 0.5; UKBB LD reference panel), generating all-but-one summary statistics spanning the entire region of the locus/super-locus. For signals in single-signal loci, we used marginal summary statistics as the all-but-one summary statistics. We repeated the conditional analysis for the male and female GWAS datasets.

### Colocalization

For each trait, we used PLINK (v.1.90b3) to calculate the LD r^2^ between all conditionally distinct GWAS lead variants and conditionally distinct adipose eQTL lead variants within 500 kb of each other using the UKBB LD reference panel described above. If the LD r^2^ was ≥ 0.5, we tested GWAS-eQTL pairs for colocalization using coloc (v.5.1.0.1, coloc.abf, default settings).^44^ For each GWAS-eQTL pair, we used the lead variant’s all-but-one eQTL and all-but-one GWAS summary statistics for the colocalization analysis. We considered GWAS-eQTL signal pairs colocalized if the coloc PP4 was ≥ 0.5. To determine the nearest gene to the GWAS signal, we used all of GENCODE v19 genes and bedtools^79^ closest function (v.2.3.0) on the GWAS lead variants. We used the same procedure to colocalize conditionally distinct signals from the female GWAS signals with female eQTL signals and to colocalize the male GWAS signals with male eQTL signals.

To compare the conditional eQTL and GWAS colocalization results to colocalization using marginal eQTL and GWAS signals, we repeated the colocalization analysis using marginal eQTL signals and GWAS full summary statistics (*P* < 5e-8). We defined the marginal eQTL signal as the most significant variant per gene in the marginal eQTL analysis for genes with ≥1 variant with *P* ≤ 1e-6.

To show visually if non-primary eQTL signals are equally likely to be colocalized compared to primary eQTL signals independent of *P*-value strength, we combined the 1^st^-5^th^ eQTL signals and divided them into four *P*-value quartiles. In each quartile and eQTL signal number we counted the number of eQTL signals and the number of eQTL signals colocalized with at least one GWAS signal. We calculated the proportion and standard error for the number of eQTL signals colocalized out of the total eQTL signals separated by signal number and quartile.

### SMR

We used SMR^10^ (v.1.3.1) on colocalized all-but-one conditionally distinct GWAS-eQTL signals using default parameters and a *P* < 1e-6 threshold to select the lead eQTL for the SMR test. Results were obtained for 3,587 of the 3,605 GWAS-eQTL signal pairs tested. We considered SMR results significant at *P* ≤ 1.4e-5 (0.05/3,605).

### MRLocus

We used MRLocus^46^ (v.0.0.26) on colocalized GWAS-eQTL signals for which multiple GWAS signals for the same trait were colocalized with multiple eQTL signals for the same gene (allelic series). To limit MRLocus analyses to nearly independent allelic series, we excluded genes for which eQTL signals had an LD r^2^ > 0.05, and then in a second analysis, we further excluded genes if eQTL signals had an LD D’ > 0.1. We also included eQTL signals that did not colocalize with GWAS signals in the MR analysis.

### Association of gene expression with metabolic traits

To follow-up the *ZNRF3*, *PDE3A*, and *SEMA3C* colocalized GWAS-eQTL pairs, we performed trait-gene expression associations in METSIM (S) using HDL-C and WHR measurements. We inverse normal transformed the gene expression and phenotypes and performed linear regression using the lm() function in R, adjusting for BMI, age, sequencing batch, RIN, mean read insert size, and read deletion size.

### Enrichment of eQTL with chromatin states and chromatin accessibility

We used GARFIELD^80^ (v2) to test for enrichment of eQTL signals in adipose tissue promoter and enhancer chromatin states from the NIH Roadmap Epigenomics project,^51^ and chromatin accessibility ATAC-seq peaks from six datasets: the top 100K peaks from SGBS cells at day 0 of differentiation (preadipocytes), day 4 (partially differentiated adipocytes), and day 14 (mature adipocytes); preadipocyte-dependent peaks; adipocyte-dependent peaks; and METSIM adipose tissue consensus peaks.^53^ We tested for enrichment separately by eQTL signal number, including the lead variants for genes without a significant eQTL as background in all analyses. We used GARFIELD to separate variants into “test” and “background” sets based on an eQTL threshold of *P* <1e-6 and estimated independent variants by clumping both the test and background variants using an LD r^2^ threshold of 0.1. We tested for overlap of both the clumped variants and their LD proxies (r^2^>0.8, PLINK, UKBB) with the regulatory elements and compared the proportion of overlaps in the test set to that in the background set for each regulatory element class using logistic regression, controlling for variant MAF, number of LD proxies, and distance to nearest gene. We used the beta from the logistic regression model, which is the natural log of the odds ratio, as the effect size and its *P*-value to assess significance. Additionally, we created a BED file of the eQTL gene, eQTL lead variants, and their proxies, and used the bedtools^79^ intersectBed function^79^ (v.2.3.0) to overlap the eQTL variants and their proxies with ATAC peak accessible chromatin regions from the same datasets as the enrichment.

### Cell culture

We cultured hWAT-A41 preadipocytes (provided by Yu-Hua Tseng, Joslin Diabetes Center^81^) in DMEM-high glucose (Sigma) supplemented with 10% fetal bovine serum (FBS). For differentiation, we plated 40,000 preadipocytes per well in a 24-well plate, grew them to confluence, and differentiated them for 5 days using induction media containing DMEM-high glucose supplemented with 2% FBS, 17 μM pantothenate, 33 μM biotin, 0.5 μM human insulin, 2 nM triiodothyronine, 0.1 μM dexamethasone, 500 μM IBMX and 30 μM indomethacin. We replaced the media every two days for five days.

We cultured SGBS preadipocytes (provided by Dr. Martin Wabitsch, University of Ulm) in basal medium (DMEM:F12, 17 μM pantothenate and 33 μM biotin) with 10% FBS. For day 5 differentiated adipocytes, we plated 40,000 preadipocytes per well in a 24-well plate, grew the cells to confluency, and induced differentiation for five days as described previously^53^.

We cultured LHCN-M2 human myoblasts (Evercyte GmbH, Vienna, Austria) as previously described^82^ in DMEM/medium 199 (Gibco, 4 +1) with 15% FBS, 0.02 M HEPES, 0.03 μg/ml zinc sulfate, 1.4 μg/ml vitamin B12, 0.055 μg/ml dexamethasone, 2.5 ng/ml recombinant human hepatocyte growth factor (Pepro Tech cat# 100-39), and 10 ng/ml basic FGF (Pepro Tech cat# 100-18B). For differentiation, we plated 25,000 LHCN-M2 myoblasts per well in a 24-well plate, grew the cells to confluency, and changed to DMEM-5.5 mM glucose with 2% horse serum for four days. We maintained all cells at 37°C in a humidified incubator with 5% CO2.

### Transcriptional reporter luciferase assay

To test the allelic differences in transcriptional activity, we designed PCR primers (**Table S25**) to amplify DNA fragments containing rs917191 (478 bp) or rs12537553 (693 bp). We generated PCR products using DNA from individuals homozygous for both alleles and cloned them into luciferase reporter vector pGL4.23 (Promega) in forward and reverse orientations with respect to the genome. We tested transcriptional activity in preadipocytes, day 3 differentiated adipocytes (hWAT and SGBS), myoblasts, and day 3 differentiated myocytes (LHCN-M2). We plated 35,000 cells per well for hWAT preadipocytes, SGBS preadipocytes, and LHCN-M2 myoblasts in 24-well plates one day before transfection. We co-transfected three sequence-verified constructs with phRL-TK Renilla reporter vector (Promega) using lipofectamine 3000 (Life Technologies) for hWAT and SGBS cells and lipofectamine LTX (Life Technologies) for LHCN-M2 cells in triplicate according to manufacturer’s protocol. We measured luciferase activity 28 hours (SGBS) or 48 hours (hWAT and LHCN-M2) post-transfection using a dual-luciferase assay system and normalized firefly luciferase activity to Renilla luciferase activity values.^83^ We quantified activity relative to an ‘empty’ vector without an added DNA fragment. For each variant, orientation, and cell type, we tested for significant (P<0.05) differences in the relative activity between the two alleles using unpaired t-tests.

### Data availability

The AdipoExpress meta-analysis results are available at https://mohlke.web.unc.edu/data/. Results include full marginal eQTL summary statistics for all ancestries, only European-ancestry individuals, males, and females, along with the conditional all-but-one eQTL summary statistics for each signal. Locus plots for every GWAS-eQTL colocalized signal pair are also available. METSIM genotypes and gene expression data are available at dbGaP phs000743.v3. FUSION genotypes and gene expression data are available at dbGaP phs001048. TwinsUK RNA-Seq data are available in the European Genome-phenome Archive (EGA) under accession EGAS00001000805. TwinsUK genotypes are available upon application to the TwinsUK Resource Executive Committee (TREC). For information on how to apply, see https://twinsuk.ac.uk/resources-for-researchers/access-our-data/.

### Code availability

All software used in this study is publicly available. Apex2R can be found here: https://github.com/corbinq/apex2R.

## Acknowledgements

We thank the METSIM, FUSION, and TwinsUK study investigators and participants for providing the subcutaneous adipose tissue samples and genotypes that made this study possible.

This study was supported by the following NIH grants: R01DK079357, R01DK072193, R01DK132775, R01DK132775, R01HL170604, R01HG010505, R01DK062370, R01HG0099765, UM1DK126185, U01DK105561, ZIAHG000024, F31HL146121, F31HL154730, T32HL129982, and T32GM007092. This study was also supported by opportunity pool funds from the Accelerating Medical Partnerships Type 2 Diabetes consortium (U01DK105554/UM1DK105554), the Academy of Finland (321428), Sigrid Juselius Foundation, Finnish Foundation for Cardiovascular Research, Centre of Excellence of Cardiovascular and Metabolic Diseases (0245896-3), Medical Research Council (MR/M004422/1 and MR/R023131/1), NIHR Biomedical Research Centre (BRC), King’s-China Scholarship Council PhD scholarship, and the Global Partnership Initiative. TwinsUK is funded by the Wellcome Trust, Medical Research Council, Versus Arthritis, European Union Horizon 2020, Chronic Disease Research Foundation (CDRF), Zoe Ltd and the National Institute for Health Research (NIHR) Clinical Research Network (CRN) and Biomedical Research Centre based at the Guy’s and St Thomas’ NHS Foundation Trust in partnership with King’s College London. This project utilized the King’s Computational Research, Engineering and Technology Environment (CREATE). This research has been conducted using the UK Biobank Resource under Application Number 25953. The Genotype-Tissue Expression (GTEx) Project was supported by the Common Fund of the Office of the Director of the National Institutes of Health (commonfund.nih.gov/GTEx). Additional funds were provided by the NCI, NHGRI, NHLBI, NIDA, NIMH, and NINDS. Donors were enrolled at Biospecimen Source Sites funded by NCI\Leidos Biomedical Research, Inc. subcontracts to the National Disease Research Interchange (10XS170), Roswell Park Cancer Institute (10XS171), and Science Care, Inc. (X10S172). The Laboratory, Data Analysis, and Coordinating Center (LDACC) was funded through a contract (HHSN268201000029C) to the The Broad Institute, Inc. Biorepository operations were funded through a Leidos Biomedical Research, Inc. subcontract to Van Andel Research Institute (10ST1035). Additional data repository and project management were provided by Leidos Biomedical Research, Inc.(HHSN261200800001E). The Brain Bank was supported supplements to University of Miami grant DA006227. Statistical Methods development grants were made to the University of Geneva (MH090941 & MH101814), the University of Chicago (MH090951,MH090937, MH101825, & MH101820), the University of North Carolina - Chapel Hill (MH090936), North Carolina State University (MH101819),Harvard University (MH090948), Stanford University (MH101782), Washington University (MH101810), and to the University of Pennsylvania (MH101822). The datasets used for the analyses described in this manuscript were obtained from dbGaP at http://www.ncbi.nlm.nih.gov/gap through dbGaP accession number phs000424.v8.p2.

## Author contributions

SMB, JSE-SM, LG, KLM, KSS, and LJS conceived and designed the study. SMB, JSE-SM, and LG generated data. SMB, JSE-SM, LG, KAB, DW, AUJ, RW, KWC, MT, SV, and MIL performed analyses. HMS, ALR, TAL, AO, LFS, NN, MRE, TY, LLB, CKR, YR, XY, SCJP, JK, PP, JT, FSC, MB, HAK, and ML provided resources. SMB, JSE-SM, LG, KLM, KSS, and LJS interpreted results and wrote the manuscript. All co-authors provided critical feedback and approved the manuscript.

## Competing interests

The authors declare no competing interests.

## Ethics declaration

The METSIM study was approved by the Ethics Committee of the University of Eastern Finland and Kuopio University Hospital in Kuopio, Finland, and written informed consent was obtained from all participants. The FUSION study was approved by the coordinating ethics committee of the Hospital District of Helsinki and Uusimaa, and written informed consent was obtained from all participants. The TwinsUK study was approved by the ethics committee at St Thomas’ Hospital London, where all the biopsies were carried out. Volunteers gave informed consent and signed an approved consent form prior to the biopsy procedure. Volunteers were supplied with an appropriate detailed information sheet regarding the research project and biopsy procedure by post prior to attending for the biopsy.

## Notes

### Competing Interest Statement

The authors have declared no competing interest.

## References

1. GTEx Consortium. The GTEx Consortium atlas of genetic regulatory effects across human tissues. Science 369, 1318–1330 (2020).

2. Umans, B. D., Battle, A. & Gilad, Y. Where are the disease-associated eQTLs? Trends Genet 37, 109–124 (2021).

3. Nicolae, D. L. et al. Trait-associated SNPs are more likely to be eQTLs: annotation to enhance discovery from GWAS. PLoS Genet. 6, e1000888 (2010).

4. Gallagher, M. D. & Chen-Plotkin, A. S. The Post-GWAS Era: From Association to Function. Am J Hum Genet 102, 717–730 (2018).

5. Nica, A. C. & Dermitzakis, E. T. Expression quantitative trait loci: present and future. Philos Trans R Soc Lond B Biol Sci 368, (2013).

6. Raulerson, C. K. et al. Adipose Tissue Gene Expression Associations Reveal Hundreds of Candidate Genes for Cardiometabolic Traits. Am. J. Hum. Genet. 105, 773–787 (2019).

7. Wu, Y. et al. Colocalization of GWAS and eQTL signals at loci with multiple signals identifies additional candidate genes for body fat distribution. Hum. Mol. Genet. 28, 4161–4172 (2019).

8. Dobbyn, A. et al. Landscape of Conditional eQTL in Dorsolateral Prefrontal Cortex and Co-localization with Schizophrenia GWAS. The American Journal of Human Genetics 102, 1169–1184 (2018).

9. Zeng, B. et al. Comprehensive Multiple eQTL Detection and Its Application to GWAS Interpretation. Genetics 212, 905–918 (2019).

10. Zhu, Z. et al. Integration of summary data from GWAS and eQTL studies predicts complex trait gene targets. Nat. Genet. 48, 481–487 (2016).

11. Hormozdiari, F. et al. Widespread Allelic Heterogeneity in Complex Traits. Am J Hum Genet 100, 789–802 (2017).

12. Jansen, R. et al. Conditional eQTL analysis reveals allelic heterogeneity of gene expression. Human Molecular Genetics 26, 1444–1451 (2017).

13. Lappalainen, T. et al. Transcriptome and genome sequencing uncovers functional variation in humans. Nature 501, 506–511 (2013).

14. Spracklen, C. N. et al. Identification of type 2 diabetes loci in 433,540 East Asian individuals. Nature 582, 240–245 (2020).

15. Zeng, B. et al. Multi-ancestry eQTL meta-analysis of human brain identifies candidate causal variants for brain-related traits. Nat Genet 54, 161–169 (2022).

16. Mostafavi, H., Spence, J. P., Naqvi, S. & Pritchard, J. K. Systematic differences in discovery of genetic effects on gene expression and complex traits. Nat Genet (2023) doi:10.1038/s41588-023-01529-1.

17. Connally, N. J. et al. The missing link between genetic association and regulatory function. eLife 11, e74970 (2022).

18. Brown, M., Greenwood, E., Zeng, B., Powell, J. E. & Gibson, G. Effect of all-but-one conditional analysis for eQTL isolation in peripheral blood. Genetics 223, iyac162 (2023).

19. Delaneau, O. et al. A complete tool set for molecular QTL discovery and analysis. Nature Communications 8, 15452 (2017).

20. Yang, J., Lee, S. H., Goddard, M. E. & Visscher, P. M. GCTA: a tool for genome-wide complex trait analysis. Am. J. Hum. Genet. 88, 76–82 (2011).

21. Quick, C. et al. A versatile toolkit for molecular QTL mapping and meta-analysis at scale. 2020.12.18.423490 Preprint at 10.1101/2020.12.18.423490 (2020).

22. GTEx Consortium. Human genomics. The Genotype-Tissue Expression (GTEx) pilot analysis: multitissue gene regulation in humans. Science 348, 648–660 (2015).

23. GTEx Consortium et al. Genetic effects on gene expression across human tissues. Nature 550, 204–213 (2017).

24. Arvanitis, M., Tayeb, K., Strober, B. J. & Battle, A. Redefining tissue specificity of genetic regulation of gene expression in the presence of allelic heterogeneity. The American Journal of Human Genetics 109, 223–239 (2022).

25. Ha Elizabeth E. & Bauer Robert C. Emerging Roles for Adipose Tissue in Cardiovascular Disease. *Arteriosclerosis*, Thrombosis, and Vascular Biology 38, e137–e144 (2018).

26. Considine, R. V. et al. Serum immunoreactive-leptin concentrations in normal-weight and obese humans. N Engl J Med 334, 292–295 (1996).

27. Cornier, M.-A. et al. The Metabolic Syndrome. Endocr Rev 29, 777–822 (2008).

28. Civelek, M. et al. Genetic Regulation of Adipose Gene Expression and Cardio-Metabolic Traits. Am. J. Hum. Genet. 100, 428–443 (2017).

29. Grundberg, E. et al. Mapping cis- and trans-regulatory effects across multiple tissues in twins. Nat Genet 44, 1084–1089 (2012).

30. El-Sayed Moustafa, J. S., et al. ACE2 expression in adipose tissue is associated with cardio-metabolic risk factors and cell type composition—implications for COVID-19. International Journal of Obesity 2022 46:8 46, 1478–1486 (2022).

31. Valencak, T. G., Osterrieder, A. & Schulz, T. J. Sex matters: The effects of biological sex on adipose tissue biology and energy metabolism. Redox Biology 12, 806–813 (2017).

32. Võsa, U. et al. Large-scale cis- and trans-eQTL analyses identify thousands of genetic loci and polygenic scores that regulate blood gene expression. Nat Genet 53, 1300–1310 (2021).

33. Karczewski, K. J. et al. The mutational constraint spectrum quantified from variation in 141,456 humans. Nature 581, 434–443 (2020).

34. Aragam, K. G. et al. Discovery and systematic characterization of risk variants and genes for coronary artery disease in over a million participants. Nat Genet 54, 1803–1815 (2022).

35. Mahajan, A. et al. Fine-mapping type 2 diabetes loci to single-variant resolution using high-density imputation and islet-specific epigenome maps. Nat Genet 50, 1505–1513 (2018).

36. Yengo, L. et al. Meta-analysis of genome-wide association studies for height and body mass index in ∼700000 individuals of European ancestry. Human Molecular Genetics 27, 3641–3649 (2018).

37. Pulit, S. L. et al. Meta-analysis of genome-wide association studies for body fat distribution in 694 649 individuals of European ancestry. Human Molecular Genetics 28, 166–174 (2019).

38. UK Biobank. Neale lab http://www.nealelab.is/uk-biobank.

39. Graham, S. E. et al. The power of genetic diversity in genome-wide association studies of lipids. Nature 600, 675–679 (2021).

40. Evangelou, E. et al. Genetic analysis of over 1 million people identifies 535 new loci associated with blood pressure traits. Nat Genet 50, 1412–1425 (2018).

41. Chen, J. et al. The trans-ancestral genomic architecture of glycemic traits. Nat Genet 53, 840–860 (2021).

42. Agrawal, S. et al. Inherited basis of visceral, abdominal subcutaneous and gluteofemoral fat depots. Nat Commun 13, 3771 (2022).

43. Wallace, C. Eliciting priors and relaxing the single causal variant assumption in colocalisation analyses. PLOS Genetics 16, e1008720 (2020).

44. Wallace, C. A more accurate method for colocalisation analysis allowing for multiple causal variants. PLOS Genetics 17, e1009440 (2021).

45. Granade, M. E. et al. Feeding desensitizes A1 adenosine receptors in adipose through FOXO1-mediated transcriptional regulation. Mol Metab 63, 101543 (2022).

46. Zhu, A. et al. MRLocus: Identifying causal genes mediating a trait through Bayesian estimation of allelic heterogeneity. PLOS Genetics 17, e1009455 (2021).

47. Jin, Y.-R. & Yoon, J. K. The R-spondin family of proteins: Emerging regulators of WNT signaling. The International Journal of Biochemistry & Cell Biology 44, 2278– 2287 (2012).

48. Tocci, J. M., Felcher, C. M., García Solá, M. E. & Kordon, E. C. R-spondin-mediated WNT signaling potentiation in mammary and breast cancer development. IUBMB Life 72, 1546–1559 (2020).

49. Nagaoka, T., Shirakawa, T., Balon, T. W., Russell, J. C. & Fujita-Yamaguchi, Y. Cyclic nucleotide phosphodiesterase 3 expression in vivo: evidence for tissue-specific expression of phosphodiesterase 3A or 3B mRNA and activity in the aorta and adipose tissue of atherosclerosis-prone insulin-resistant rats. Diabetes 47, 1135– 1144 (1998).

50. Hanna, R. et al. Cardiac Phosphodiesterases Are Differentially Increased in Diabetic Cardiomyopathy. Life Sciences 283, 119857 (2021).

51. Kundaje, A. et al. Integrative analysis of 111 reference human epigenomes. Nature 518, 317–330 (2015).

52. Fischer-Posovszky, P., Newell, F. S., Wabitsch, M. & Tornqvist, H. E. Human SGBS Cells – a Unique Tool for Studies of Human Fat Cell Biology. Obes Facts 1, 184–189 (2008).

53. Perrin, H. J. et al. Chromatin accessibility and gene expression during adipocyte differentiation identify context-dependent effects at cardiometabolic GWAS loci. PLOS Genetics 17, e1009865 (2021).

54. Mejhert, N. et al. Semaphorin 3C is a novel adipokine linked to extracellular matrix composition. Diabetologia 56, 1792–1801 (2013).

55. Scott, L. J. et al. The genetic regulatory signature of type 2 diabetes in human skeletal muscle. Nature Communications 7, 11764 (2016).

56. Laakso, M. et al. METabolic Syndrome In Men (METSIM) Study: a resource for studies of metabolic and cardiovascular diseases. J. Lipid Res. 58(3), 481–493 (2017).

57. Brotman, S. M., Oravilahti, A., Rosen, J. D., Alvarez, M. & Heinonen, S. Cell-type composition affects adipose gene expression associations with cardiometabolic traits. Diabetes.

58. Hannon, G. J. FASTX-Toolkit. FASTX-Toolkit.

59. Martin, M. Cutadapt removes adapter sequences from high-throughput sequencing reads. EMBnet.journal 17, 10–12 (2011).

60. Dobin, A. et al. STAR: ultrafast universal RNA-seq aligner. Bioinformatics 29, 15– 21 (2013).

61. Hysi, P. G. et al. A genome-wide association study for myopia and refractive error identifies a susceptibility locus at 15q25. Nature Genetics 42, 902–905 (2010).

62. Buil, A. et al. Gene-gene and gene-environment interactions detected by transcriptome sequence analysis in twins. Nature Genetics 47, 88–91 (2015).

63. Glastonbury, C. A. A. et al. Adiposity-Dependent Regulatory Effects on Multi-tissue Transcriptomes. American Journal of Human Genetics 99, 567–579 (2016).

64. McCarthy, S. et al. A reference panel of 64,976 haplotypes for genotype imputation. Nature Genetics 48, 1279–1283 (2016).

65. Frankish, A. et al. GENCODE reference annotation for the human and mouse genomes. Nucleic Acids Research 47, D766–D773 (2019).

66. Hartley, S. W. & Mullikin, J. C. QoRTs: a comprehensive toolset for quality control and data processing of RNA-Seq experiments. BMC Bioinformatics 16, 224 (2015).

67. Robinson, M. D. & Oshlack, A. A scaling normalization method for differential expression analysis of RNA-seq data. Genome Biology 11, R25 (2010).

68. Robinson, M. D., McCarthy, D. J. & Smyth, G. K. edgeR: a Bioconductor package for differential expression analysis of digital gene expression data. Bioinformatics 26, 139–140 (2010).

69. Stegle, O., Parts, L., Piipari, M., Winn, J. & Durbin, R. Using probabilistic estimation of expression residuals (PEER) to obtain increased power and interpretability of gene expression analyses. Nat Protoc 7, 500–507 (2012).

70. Bates, D., Mächler, M., Bolker, B. & Walker, S. Fitting Linear Mixed-Effects Models Using lme4. Journal of Statistical Software 67, 1–48 (2015).

71. Ongen, H., Buil, A., Brown, A. A., Dermitzakis, E. T. & Delaneau, O. Fast and efficient QTL mapper for thousands of molecular phenotypes. Bioinformatics 32, 1479–1485 (2016).

72. Quick, C. apex2R. (2021).

73. Chang, C. C. et al. Second-generation PLINK: rising to the challenge of larger and richer datasets. GigaScience 4, s13742–015-0047–8 (2015).

74. Bycroft, C. et al. The UK Biobank resource with deep phenotyping and genomic data. Nature 562, 203–209 (2018).

75. Wickham, H. ggplot2: Elegant Graphics for Data Analysis. (Springer-Verlag New York, 2016).

76. Willer, C. J., Li, Y. & Abecasis, G. R. METAL: fast and efficient meta-analysis of genomewide association scans. Bioinformatics 26, 2190–2191 (2010).

77. Brown, G. W. & Mood, A. M. On Median Tests for Linear Hypotheses. Proceedings of the Second Berkeley Symposium on Mathematical Statistics and Probability 2, 159–167 (1951).

78. Pruim, R. J. et al. LocusZoom: regional visualization of genome-wide association scan results. Bioinformatics 26, 2336–2337 (2010).

79. Quinlan, A. R. & Hall, I. M. BEDTools: a flexible suite of utilities for comparing genomic features. Bioinformatics 26, 841–842 (2010).

80. Iotchkova, V. et al. GARFIELD classifies disease-relevant genomic features through integration of functional annotations with association signals. Nat Genet 51, 343–353 (2019).

81. Shamsi, F. & Tseng, Y.-H. Protocols for generation of immortalized human brown and white preadipocyte cell lines. Methods Mol Biol 1566, 77–85 (2017).

82. Zhu, C.-H. et al. Cellular senescence in human myoblasts is overcome by human telomerase reverse transcriptase and cyclin-dependent kinase 4: consequences in aging muscle and therapeutic strategies for muscular dystrophies. Aging Cell 6, 515– 523 (2007).

83. Fogarty, M. P., Cannon, M. E., Vadlamudi, S., Gaulton, K. J. & Mohlke, K. L. Identification of a Regulatory Variant That Binds FOXA1 and FOXA2 at the CDC123/CAMK1D Type 2 Diabetes GWAS Locus. PLOS Genetics 10, e1004633 (2014).

