## Supplemental Figures for "Adipose tissue eQTL meta-analysis reveals the contribution of allelic heterogeneity to gene expression regulation and cardiometabolic traits"

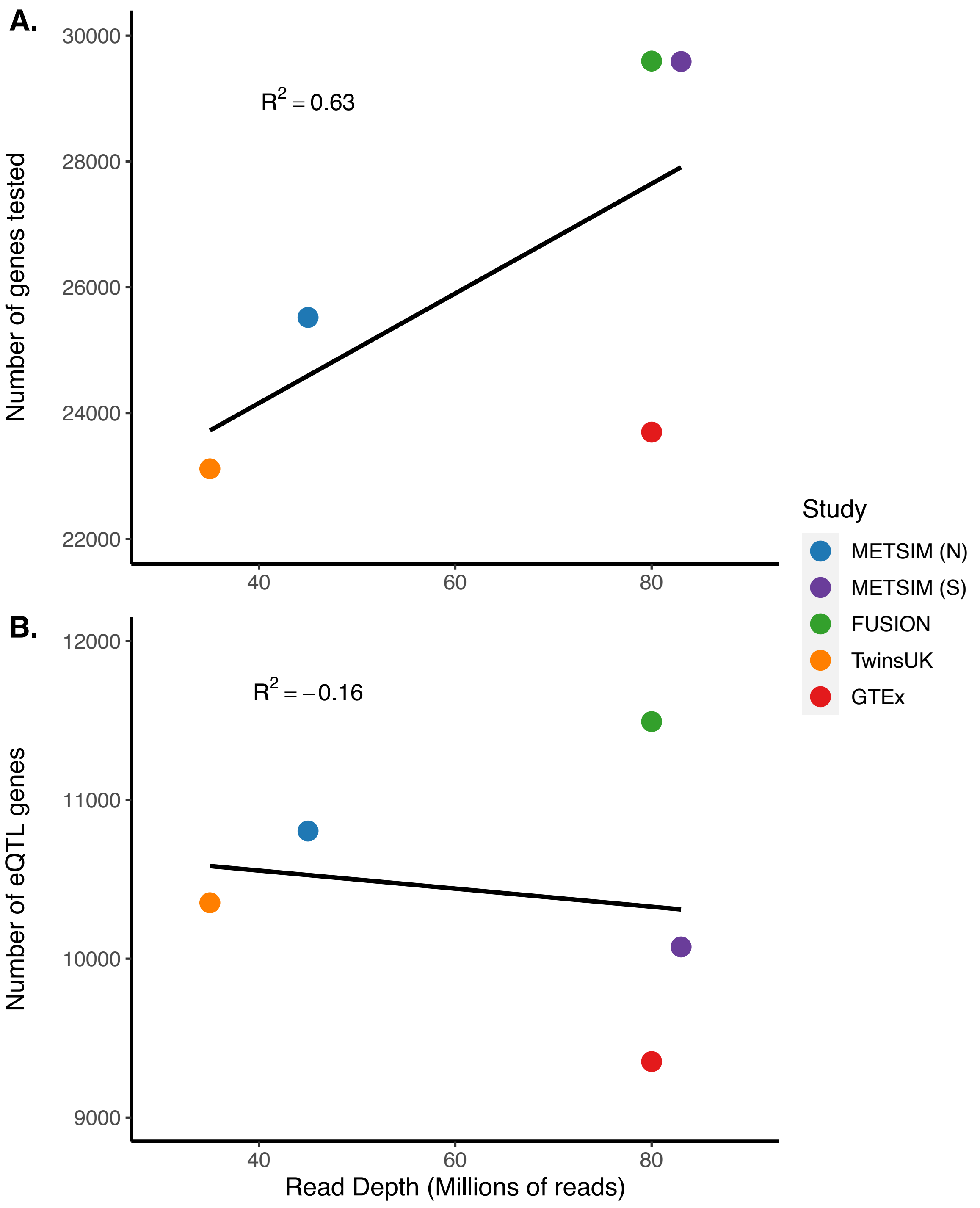


**Figure S1. Comparison of read depth and the number of genes tested in four studies.** The read depth (x-axis) versus (A) the number of genes tested for an eQTL and (B) the number of eQTL genes in all five studies. The linear regression line is shown in black and the Pearson correlation is shown by the $R^2$ on the plot.

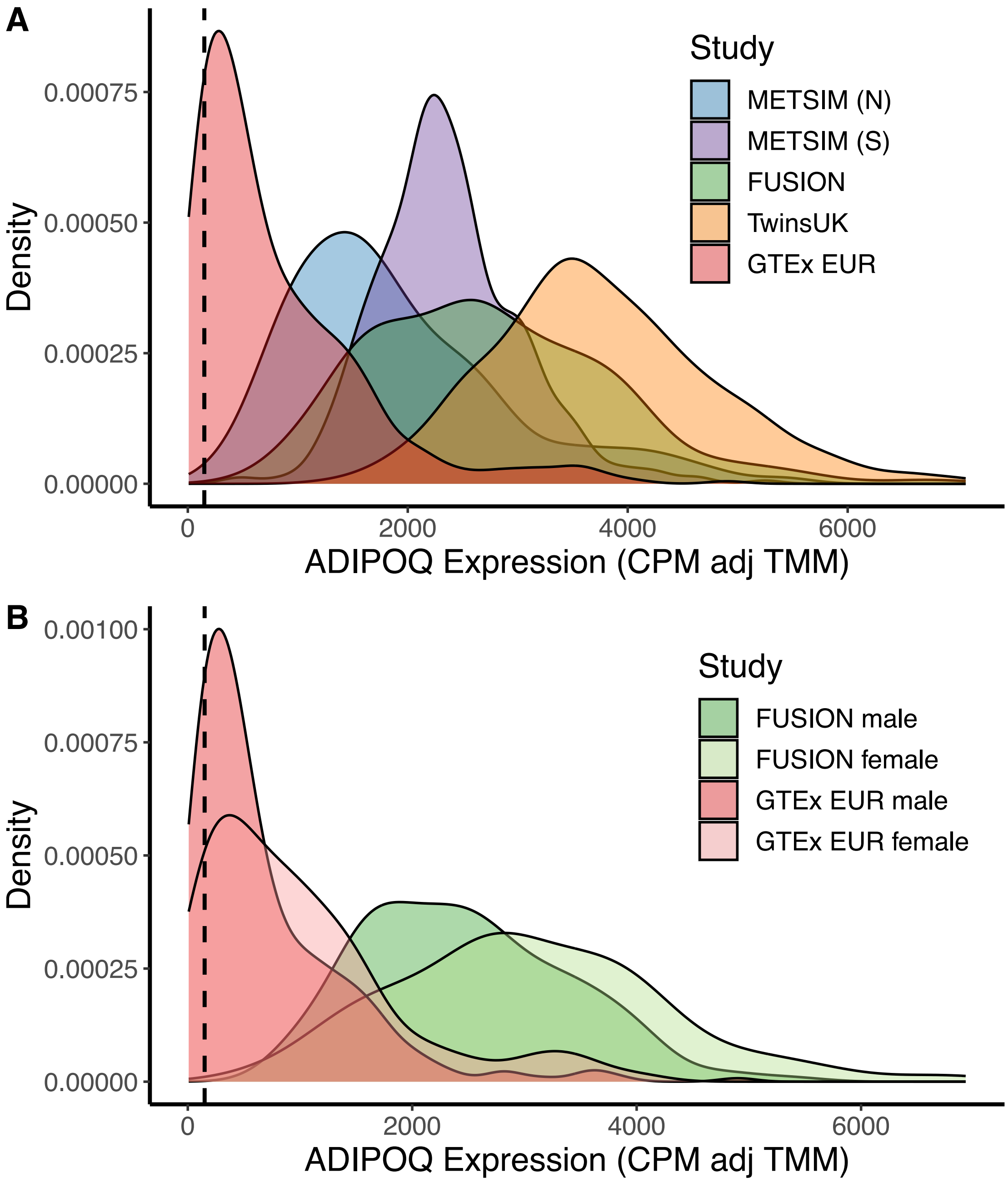


**Figure S2. Distribution of *ADIPOQ* expression for each study.**
Density plot of *ADIPOQ* expression in each study for (A) all studies and (B) FUSION and GTEx separated by sex. The x-axis is the counts per million normalized for trimmed mean of M values. The y-axis is the density. Each study is represented by a different color as indicated by the legend. The vertical black dashed line is at 150 CPM which is the threshold for retaining samples in eQTL analyses.

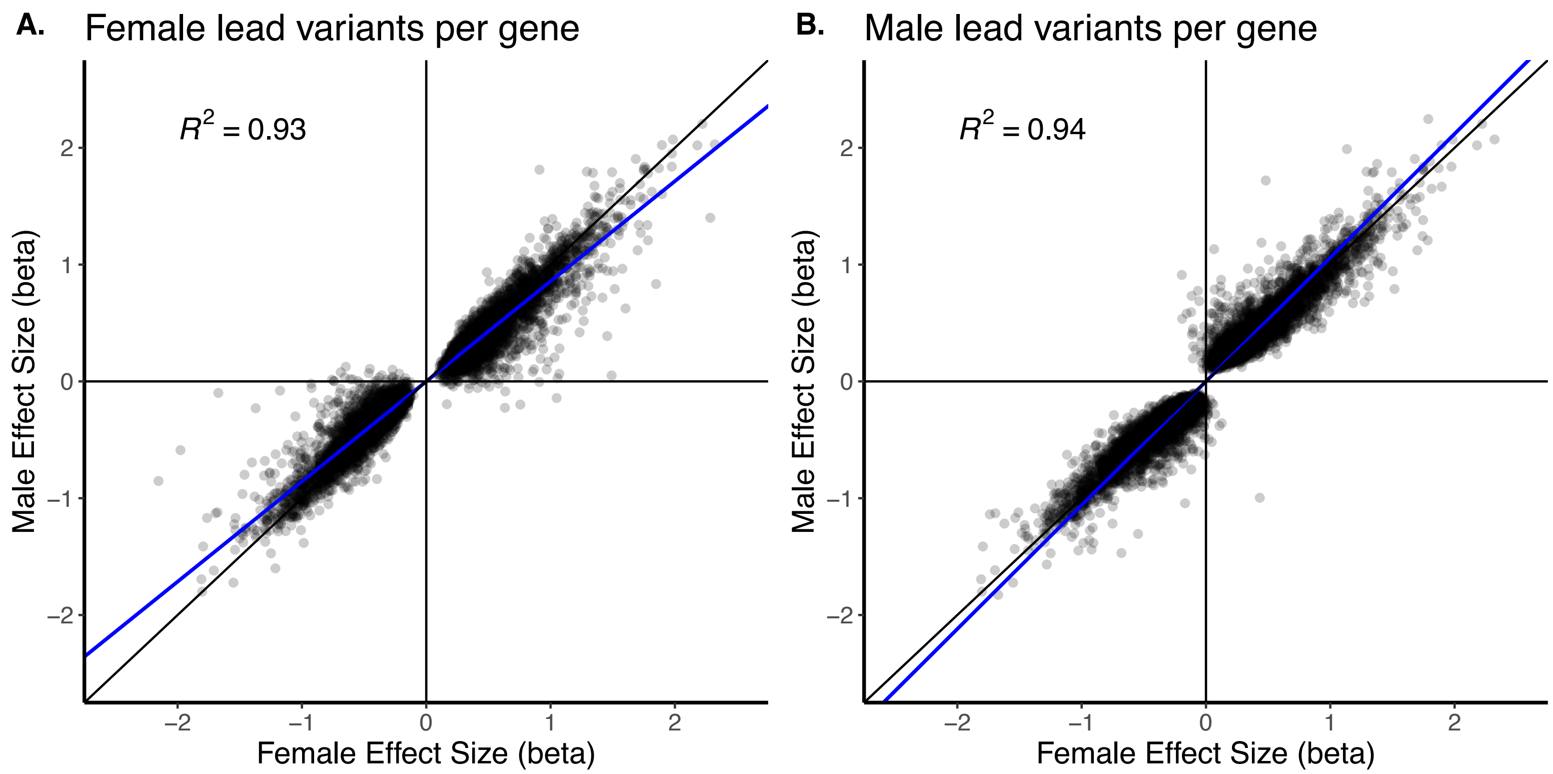


**Figure S3. Comparison of effect sizes in sex-stratified meta-analyses.**
(A) Effect sizes of lead variants per gene identified in the female marginal eQTL meta-analysis looked up in the male marginal eQTL analysis. (B) Effect sizes of lead variants per gene identified in the male marginal eQTL meta-analysis looked up in the female marginal eQTL analysis. Each point is a variant-gene pair. The linear regression lines are shown in blue and the black diagonal line indicates slope = 1. $R^2$ values are the Pearson correlations.

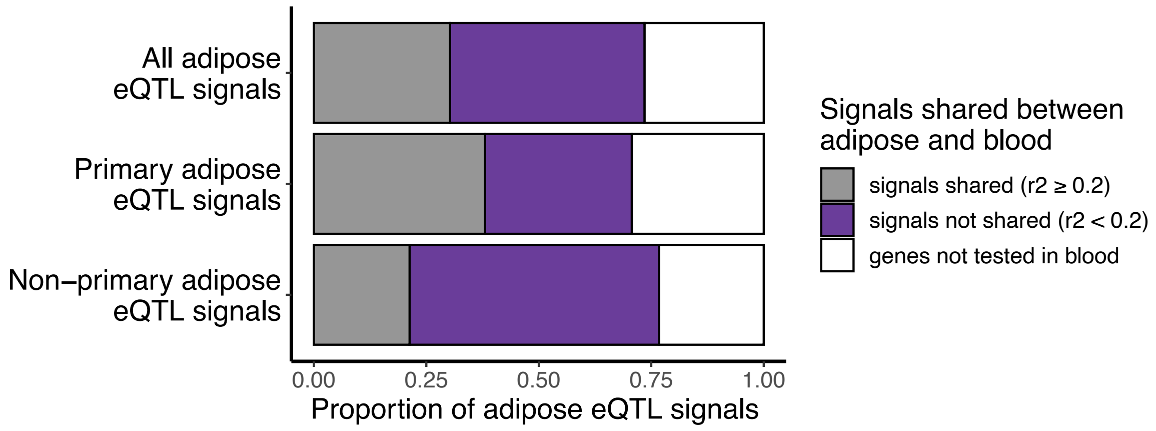


**Figure S4. Overlap of adipose and blood eQTL signals.**
Horizontal stacked bar charts show the proportion of significant adipose eQTL signals that overlap with significant blood eQTL signals (FDR ≤ 0.05). The top bar shows all adipose eQTL signals, the middle bar shows only primary adipose eQTL signals, and the bottom bar shows non-primary adipose eQTL signals. The gray indicates the proportion of adipose eQTL signals with a shared blood eQTL (LD $r^2 \geq 0.2$), the purple indicates the proportion of adipose eQTL signals not shared with a blood eQTL (LD $r^2 < 0.2$), and white indicates the eQTL genes in adipose not tested in blood.

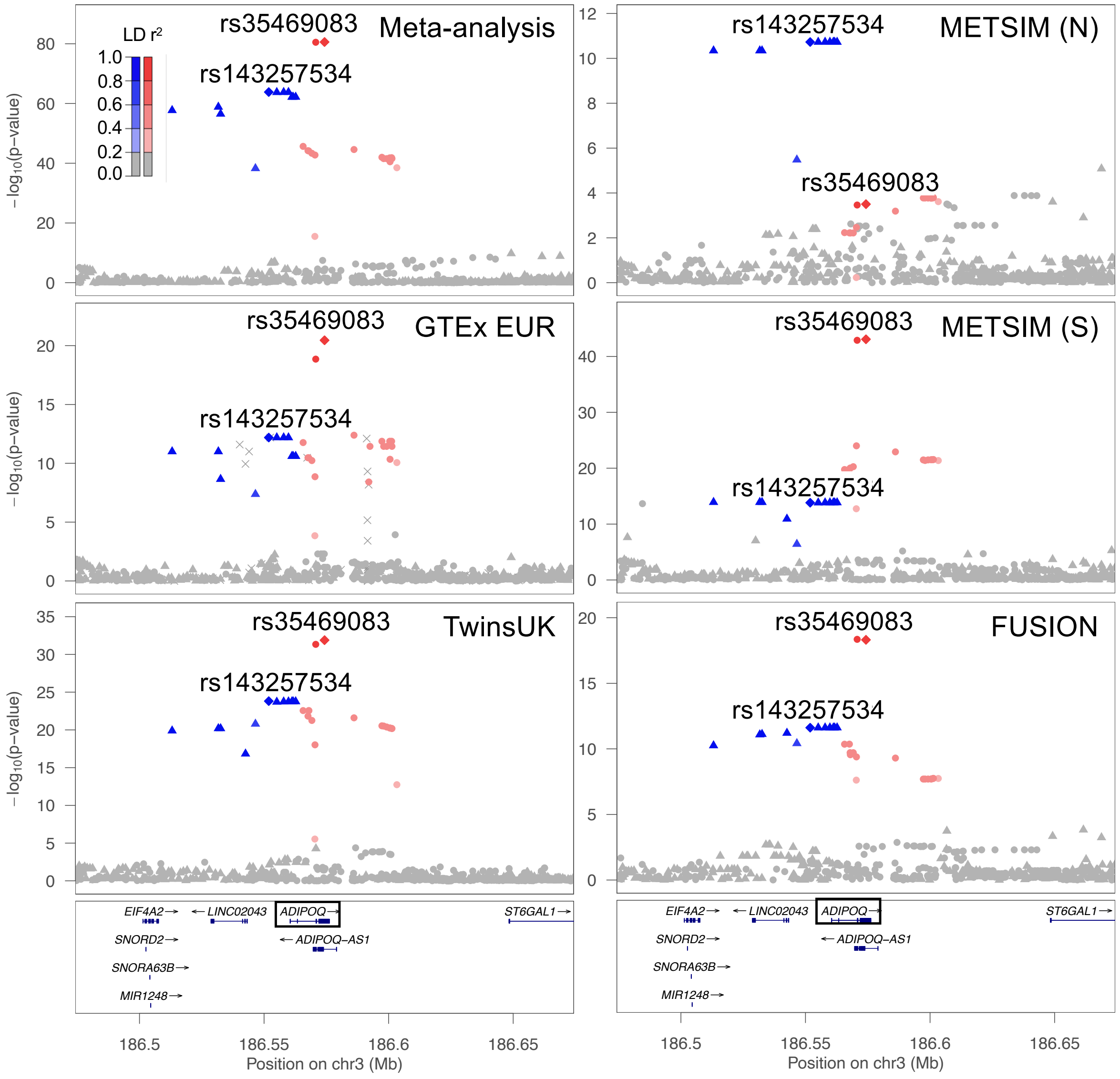


**Figure S5. *ADIPOQ* eQTL signals in each study versus the meta-analysis.**
LocusZoom plots of the marginal *ADIPOQ* eQTL for the meta-analysis and each individual study. The x-axes show position on chromosome 3 and y-axes show eQTL $-\log_{10}$ *P*-value. The lead variant of the 1st signal (chr3:186,574,282, rs35469083) in the meta-analysis is represented by a red diamond in all plots, and the lead variant of the 2nd signal (chr3:186,551,888, rs143257534) in the meta-analysis is represented by a blue diamond in all plots. The red circles represent variants in stronger LD with the lead variant of the 1st signal while the blue triangles represent variants in stronger LD with the lead variant of the 2nd signal. Shading indicates LD $r^2$ as shown in the legend. Although each study has both signals colored, only one signal was significant in the conditional eQTL analysis for each of the individual studies ($P \leq$ 1e-6).

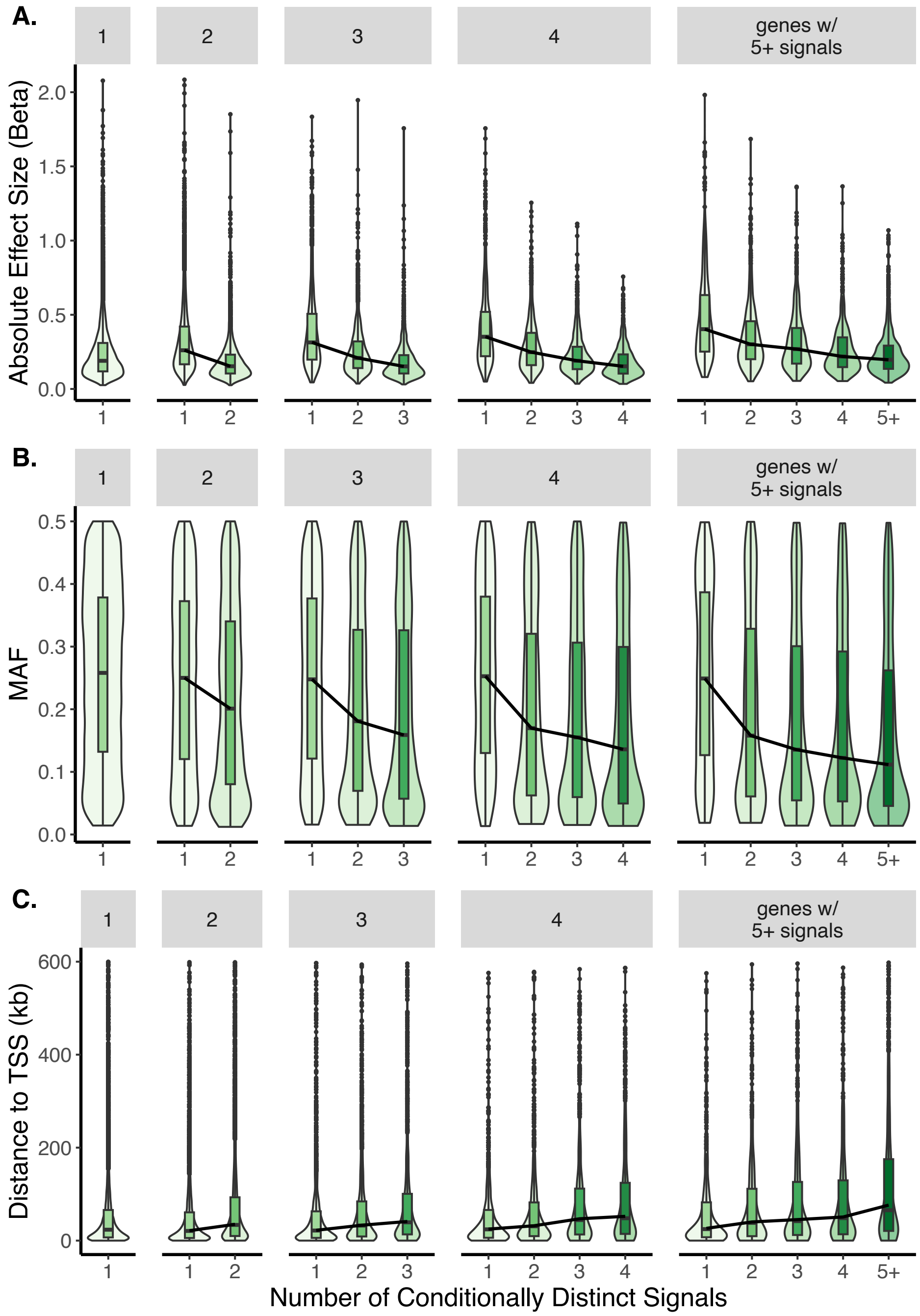


**Figure S6. Characteristics of eQTL signal variants.**
Violin plots with inset boxplots of the (A) absolute value of the effect sizes of lead variants, (B) MAF, and (C) distance of the lead variants to the gene TSS for the indicated signals in order of discovery. Each plot represents 5 subgroups which are genes with only one signal, genes with exactly 2 signals, genes with exactly 3 signals, genes with exactly 4 signals, and genes with 5 or more signals. For the boxplots, the center line represents the median value, the box limits represent the upper and lower quartiles, whiskers represent the 1.5x interquartile range, and the black circles represent outliers. The black lines connect the median values of each signal group. In C, 163 points with a distance to TSS greater than 600 were excluded.

A.

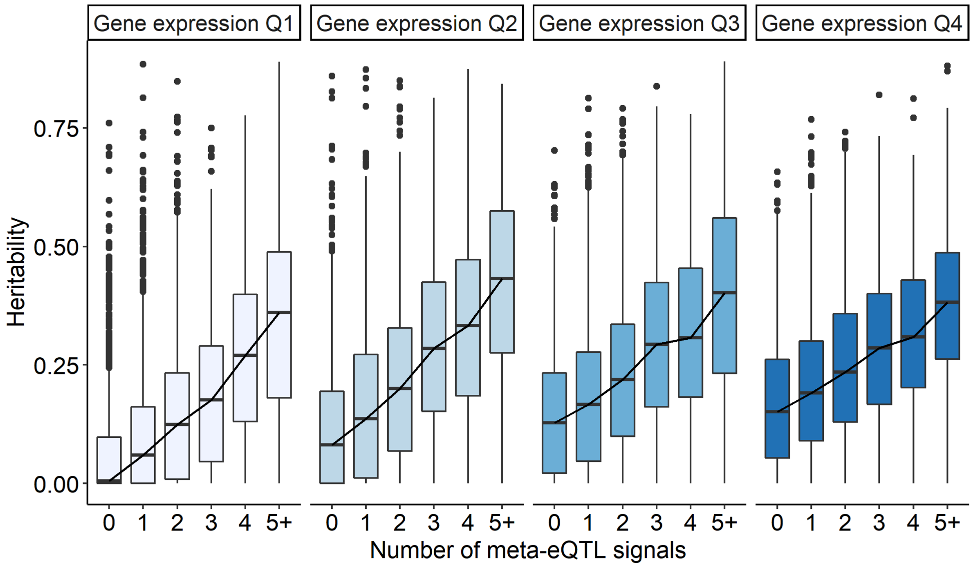


B.

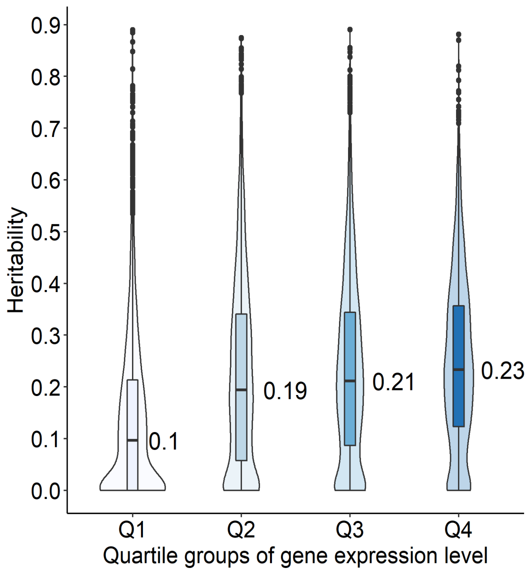


**Figure S7. Heritability of genes with an eQTL grouped by gene expression.**
(A) Boxplots of heritability for genes with zero to five or more eQTL signals. Boxplots are split into subgroups based on the quartile of expression for the gene. Lowly expressed genes are in quartile 1 shown in white while highly expressed genes are in quartile 4 shown in darkest blue. The center line represents the median value, the box limits represent the upper and lower quartiles, whiskers represent the 1.5x interquartile range, and the black circles represent outliers. The black line connects the median of the boxplots. (B) Boxplots of heritability for genes separated by gene expression quartile. The number shown represents the median heritability value.

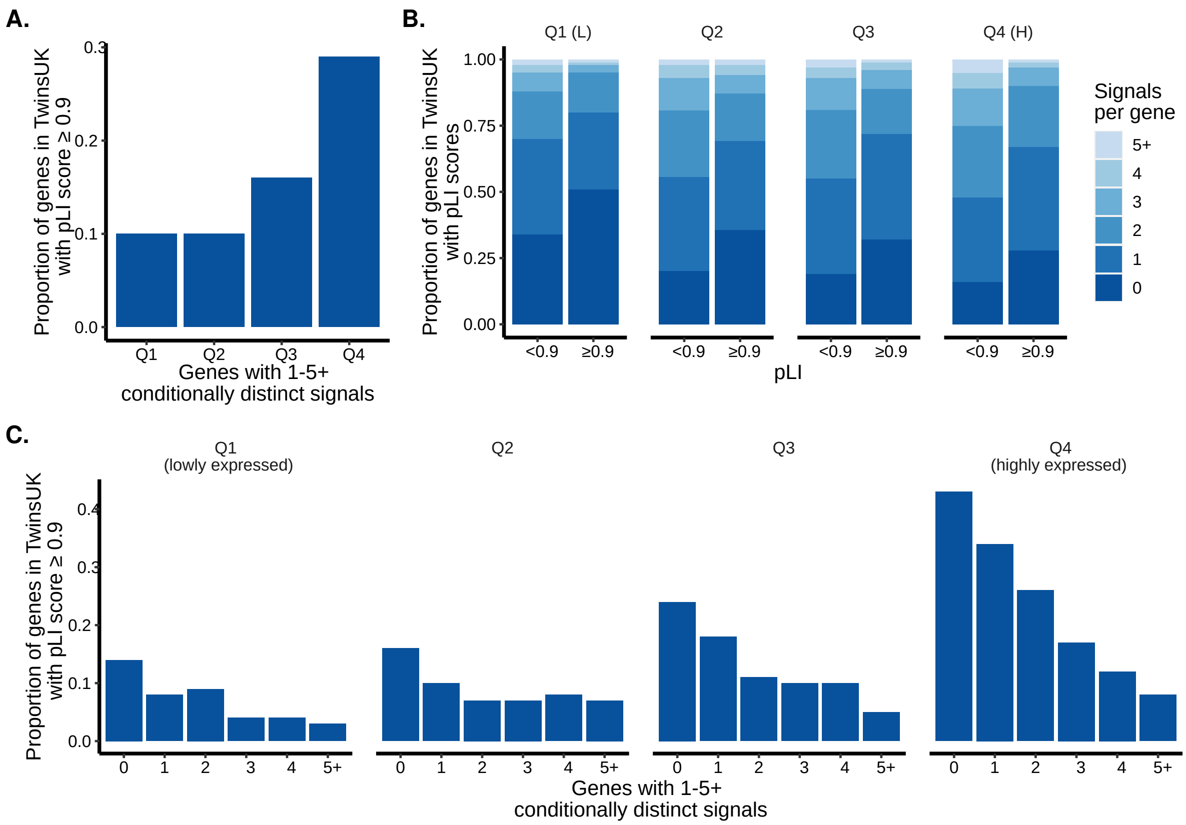


**Figure S8. pLI scores of eQTL genes with 1 through 5 or more signals.**
(A) Proportion of genes in TwinsUK for each gene expression quartile with a pLI score ≥ 0.9. Quartile 1 indicates the genes with the lowest expression and quartile 4 indicates genes with the highest expression. (B) Proportion of genes in TwinsUK with a pLI score < 0.9 or ≥ 0.9 separated by signal number for each gene expression quartile. The darkest blue are the genes without an eQTL signal and the lightest blue are genes with five or more eQTL signals. For each expression level quartile, the proportion of genes with multiple signals was substantially lower for genes with pLI ≥0.9 than for genes with pLI <0.9: this trend was particularly pronounced in the highest expression category. (C) Proportion of genes in TwinsUK for each eQTL signal number with a pLI score ≥ 0.9. Each set of bar graphs is for a different gene expression quartile.

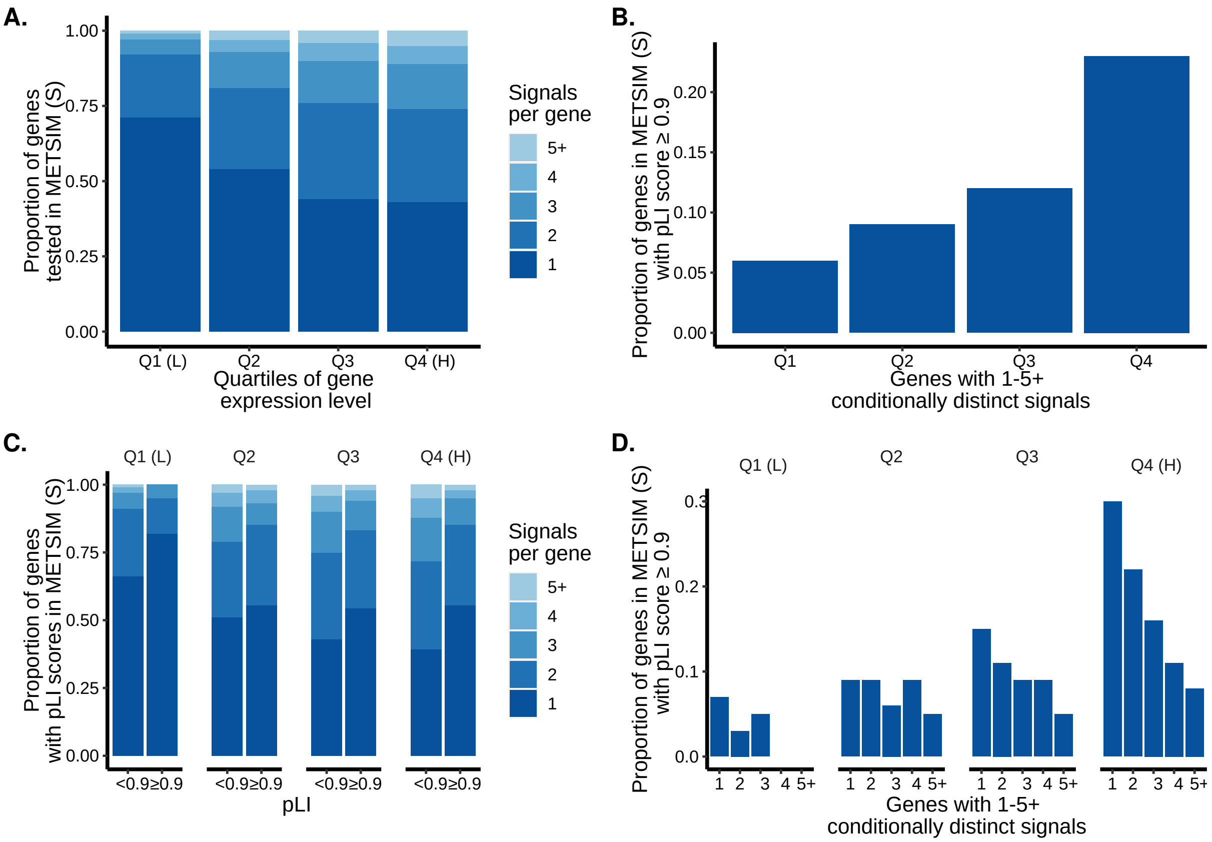


**Figure S9. Gene expression levels in METSIM surgical biopsies and pLI scores of eQTL genes with 1 through 5 or more signals.**

(A) Proportion of genes with the specified number of eQTL signals separated by gene expression quartiles. Quartile 1 indicates the genes with the lowest expression. The darkest blue are the genes without an eQTL signal and the lightest blue are genes with five or more eQTL signals. (B) Proportion of genes for each gene expression quartile with a pLI score ≥ 0.9. Quartile 1 indicates the genes with the lowest expression. (C) Proportion of genes with a pLI score ≥ 0.9 separated by signal number for each gene expression quartile. The darkest blue are the genes without an eQTL signal and the lightest blue are genes with five or more eQTL signals. (D) Proportion of genes for each eQTL signal number with a pLI score ≥ 0.9. Each set of bar graphs is for a different gene expression quartile.

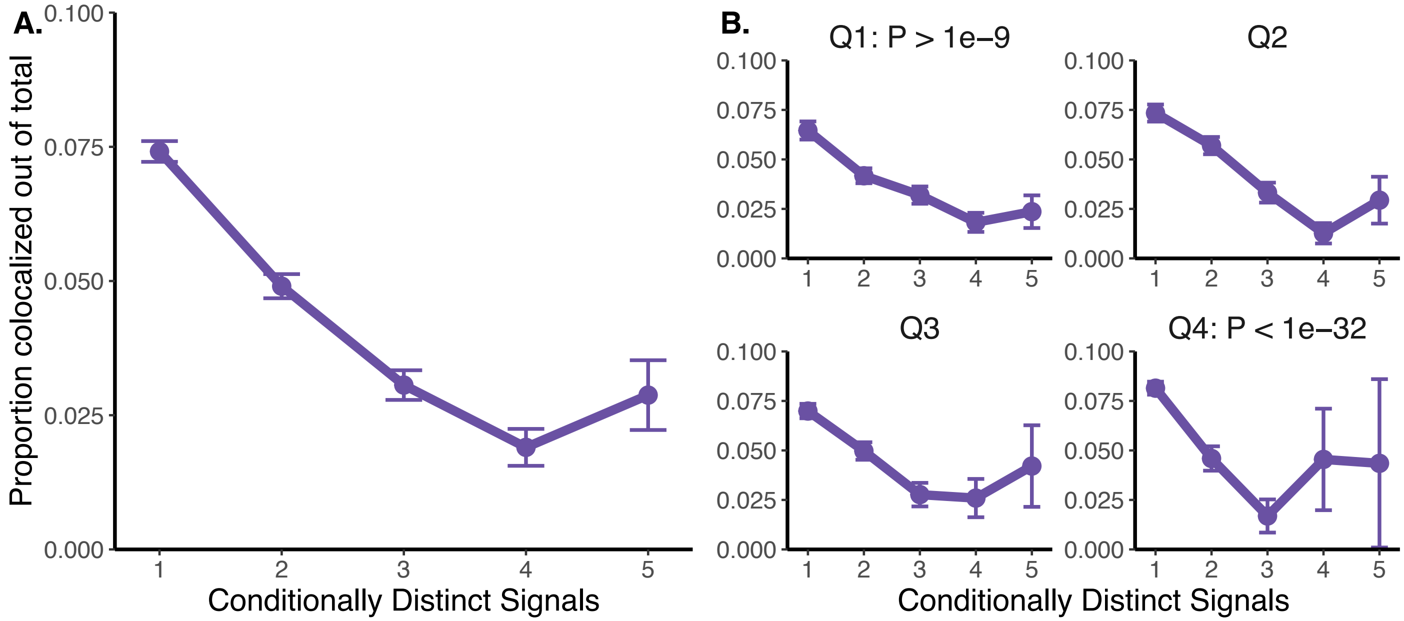


**Figure S10. Proportion of eQTL signals colocalized with GWAS signals.**
Proportion of eQTL signals colocalized with a GWAS signal out of the total number of eQTL signals separated by signal number. (A) Includes all eQTL signals and (B) separates the eQTL signals by quartiles of *P*-value strength. The top left are the least significant eQTL signals, the top right are the second least significant eQTL signals, the bottom left are the second most significant eQTL signals, and the bottom right are the most significant eQTL signals. The standard error is shown by the error bars.

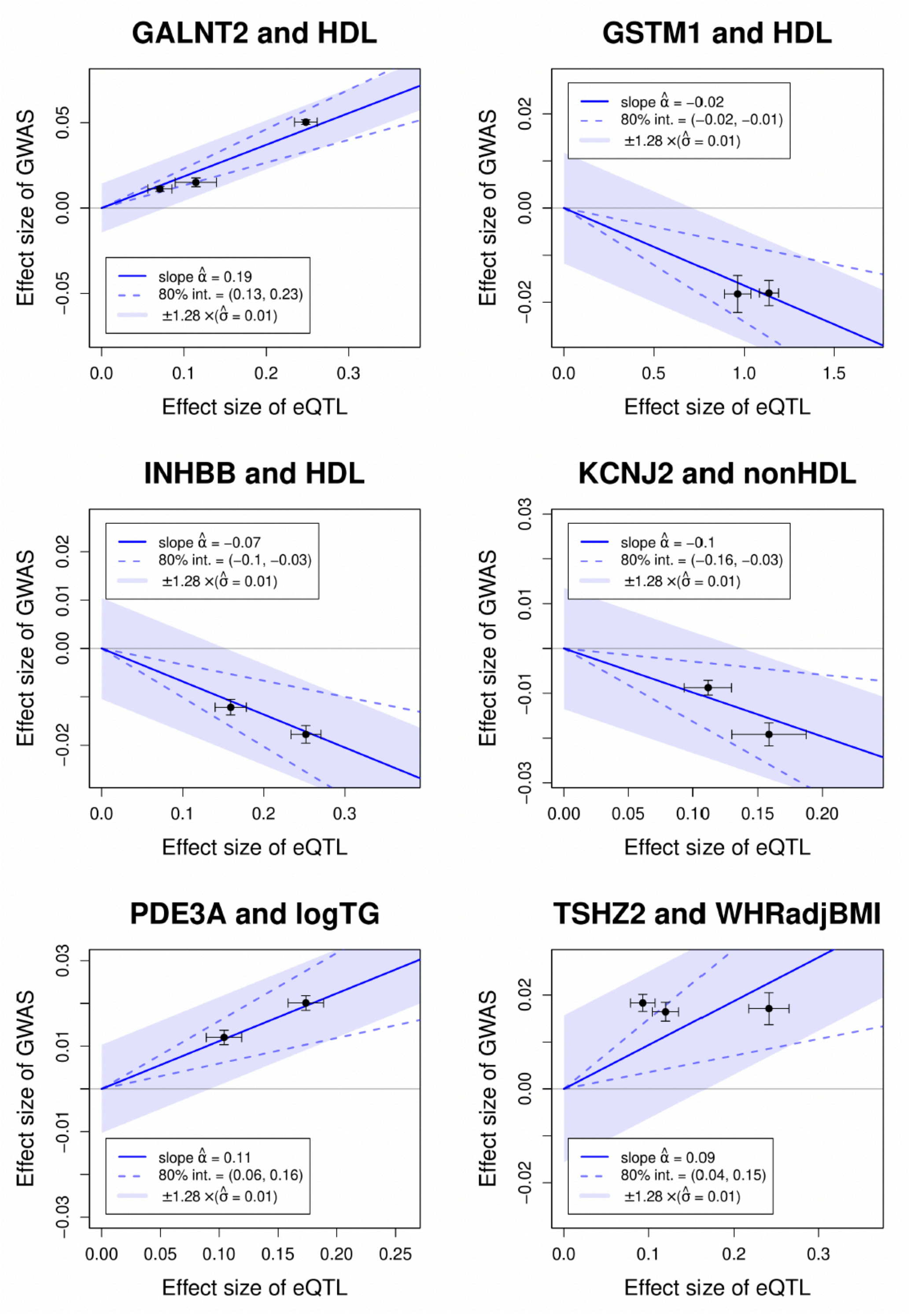


**Figure S11. Mendelian randomization using MRLocus for select allelic series.**
Plots of the effect size of the GWAS signals (y-axis) versus the effect size of the eQTL signals (x-axis) from MRLocus. Six eQTL gene-trait pairs are included with at least 2 signals colocalized with each other. Each point represents a colocalized eQTL signal with standard error bars. The solid blue lines represent the overall slope of the effect of gene on the trait, and dotted blue lines represent the confidence intervals.

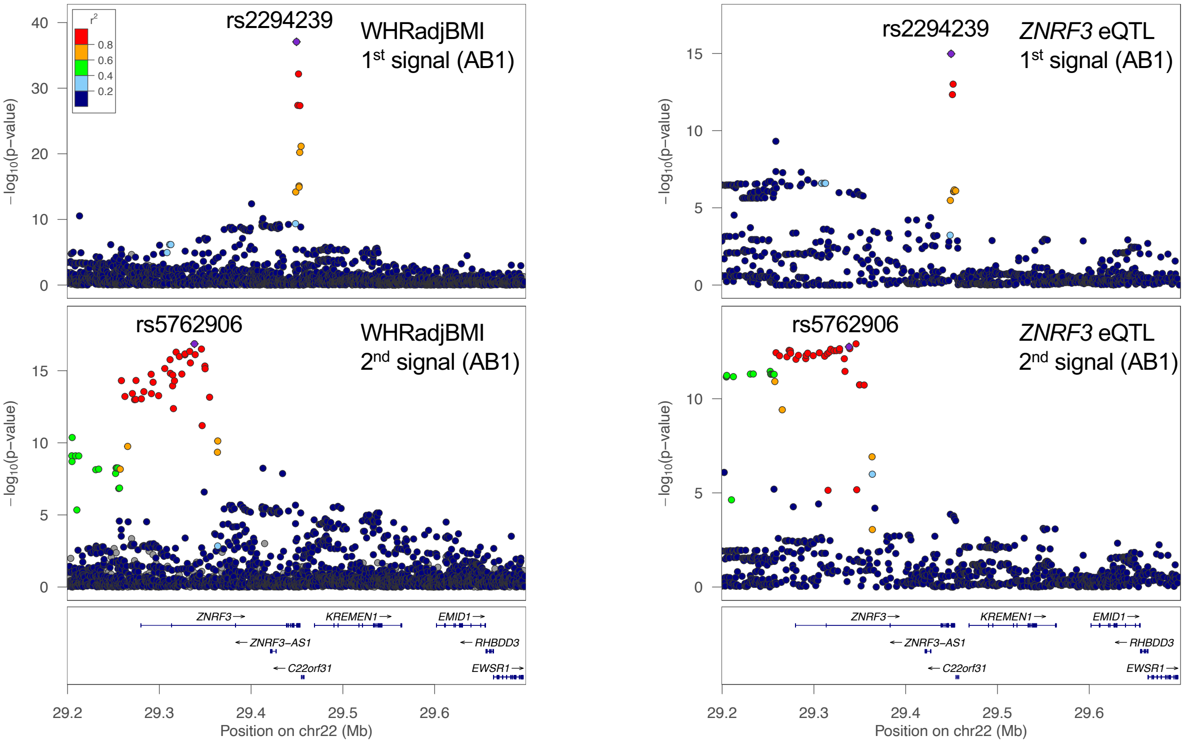


**Figure S12. WHRadjBMI GWAS and *ZNRF3* eQTL conditional signal plots.** LocusZoom plots for WHRadjBMI signal 1 conditioned on signal 2 (top left) and WHRadjBMI signal 2 conditioned on signal 1 (bottom left). Plots for *ZNRF3* signal 1 conditioned on signal 2 (top right) and *ZNRF3* signal 2 conditioned on signal 1 (bottom right). The dots are colored by LD with the lead variant. Both plots are colored by the GWAS lead variant represented by a purple diamond.

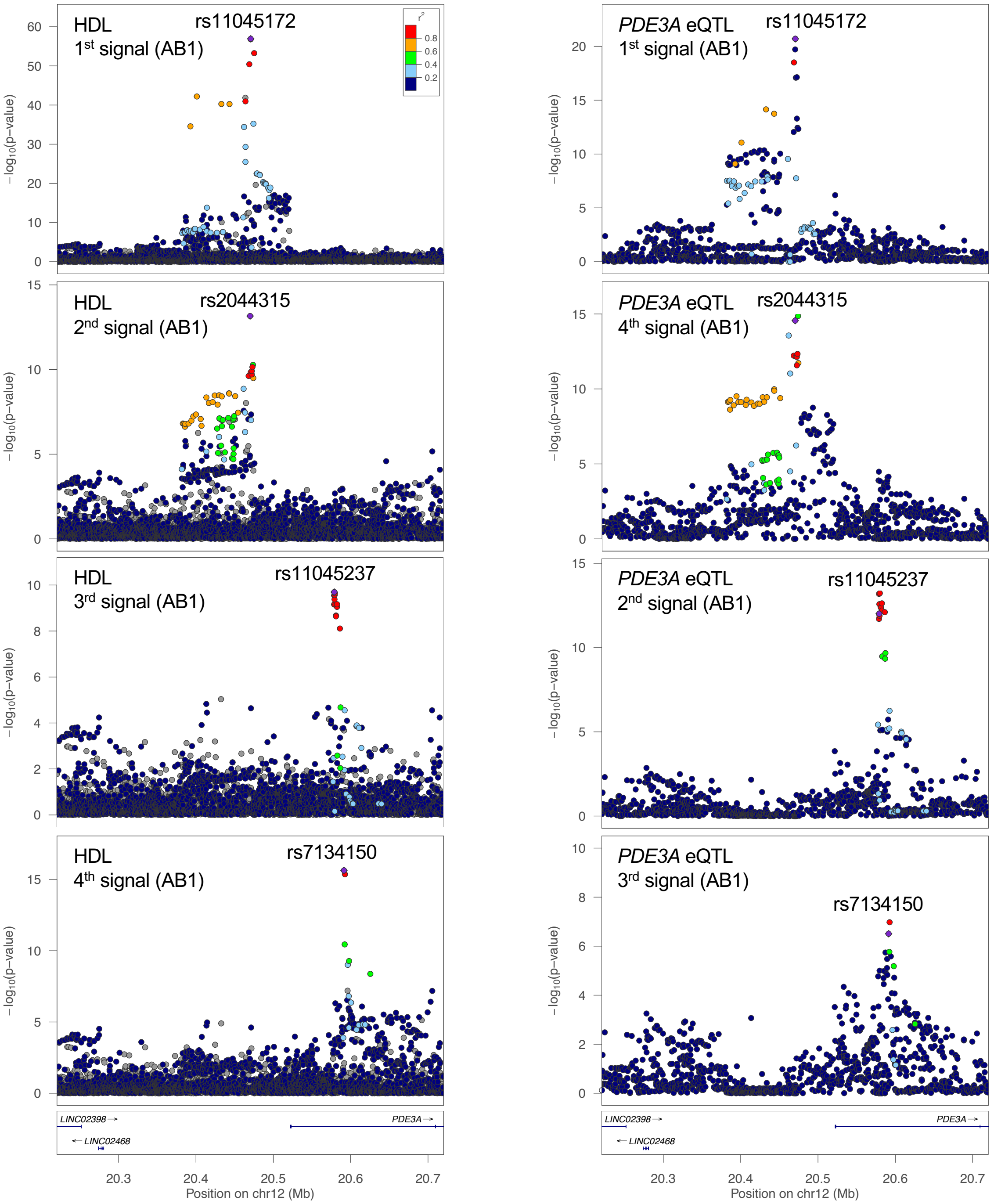


**Figure S13. HDL-C GWAS and *PDE3A* eQTL conditional signal plots.**
LocusZoom plot for HDL-C signal 1 conditioned on all the other signals, followed by plots for each signal conditioned on all other signals (left). Plot for *PDE3A* signal 1 conditioned on all the other signals, followed by plots for signal 4 conditioned on all other signals, then signal 2 and signal 3 (right). y-axes show the -$\log_{10}$ *P*-value after conditioning. The dots are colored by LD with the lead variant. Both plots are colored by the GWAS lead variant represented by a purple diamond.

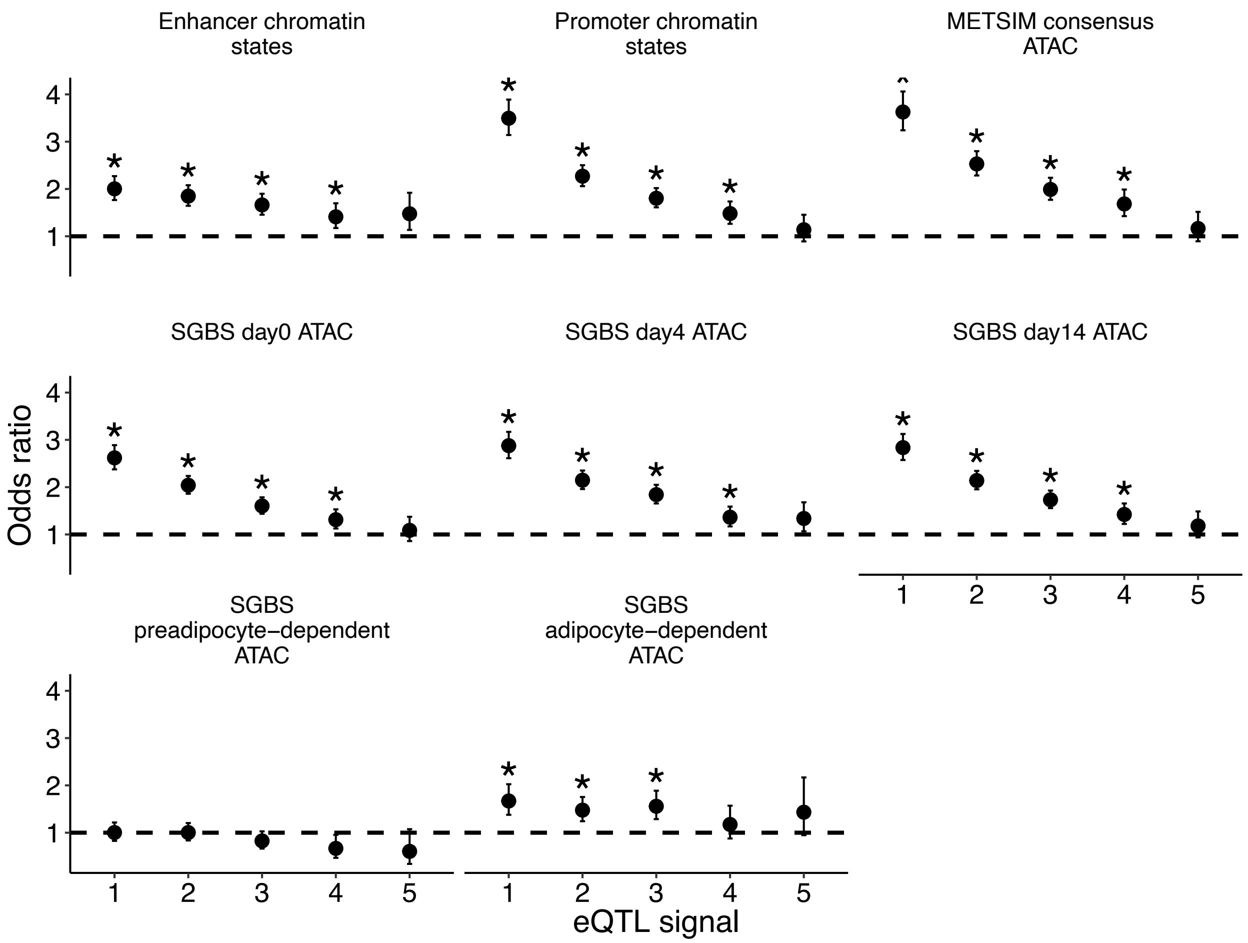


**Figure S14. eQTL signals enriched in chromatin states and ATAC-seq peaks.**
eQTL signals enriched in Roadmap Epigenomics adipose chromatin states and ATAC-seq peaks in adipose tissue, preadipocytes, immature adipocytes, and adipocytes separated by signal number. Only signals 1-5 were included. The x-axes indicate the eQTL signal number while the y-axes indicate the odds ratio (OR) for the enrichment. The bars represent the upper and lower 95% confidence intervals. Asterisks represent significant Bonferroni-adjusted enrichment values that do not overlap an OR of 1 (black dashed lines).

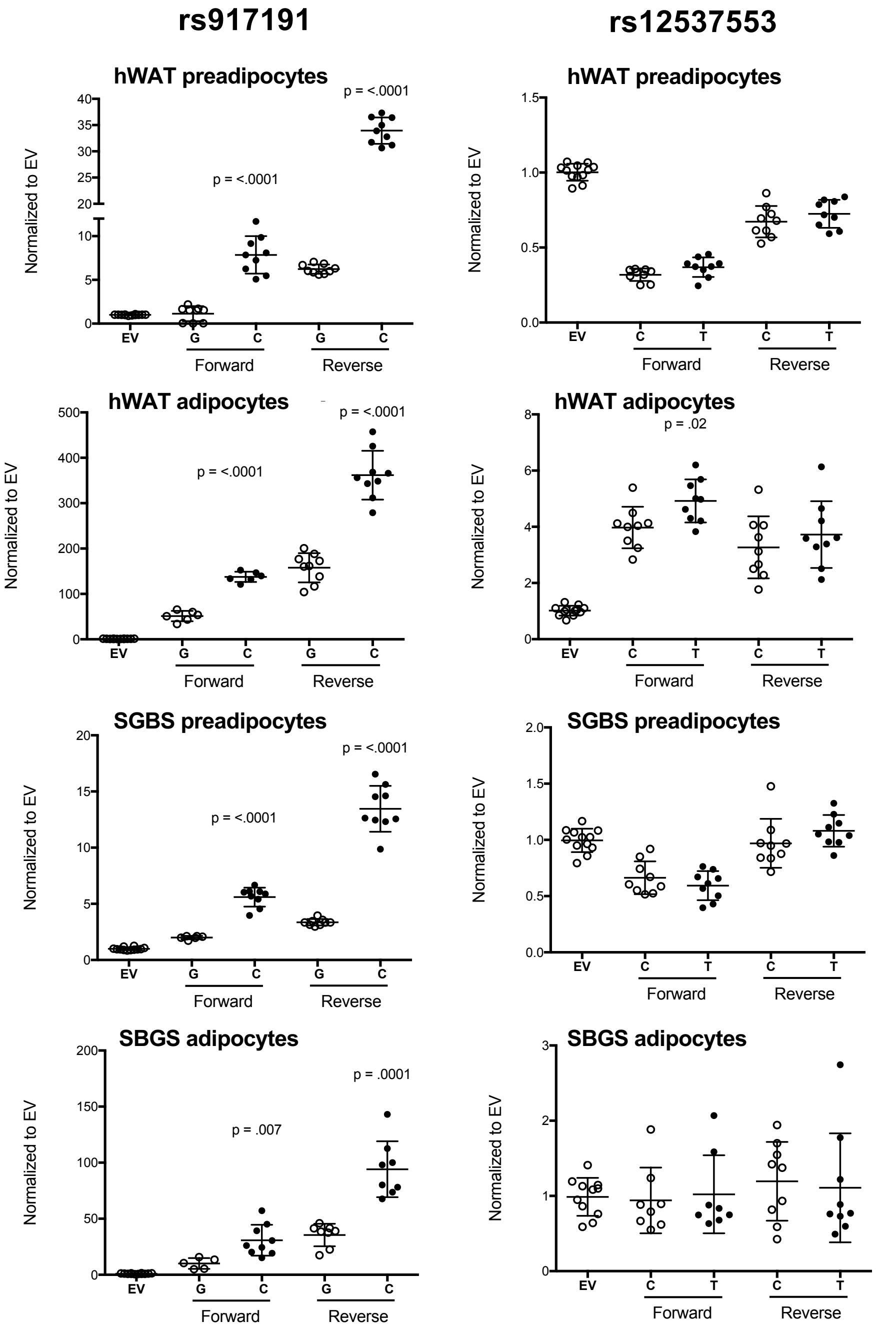


**Figure S15. Transcriptional reporter assays in preadipocytes and adipocytes for two WHRadjBMI candidate variants near *SEMA3C*.**

Transcriptional reporter assays testing activity of rs917191 (left) and rs12537553 (right) alleles in hWAT preadipocytes, hWAT differentiated adipocytes, SGBS preadipocytes, and SGBS differentiated adipocytes. Points indicate relative luciferase activity of independent clones compared to empty vector and lines represent standard error bars. *P*-values are for Student's unpaired t-tests to compare alleles. rs917191 shows strong effects and significant allelic differences in transcriptional activity.

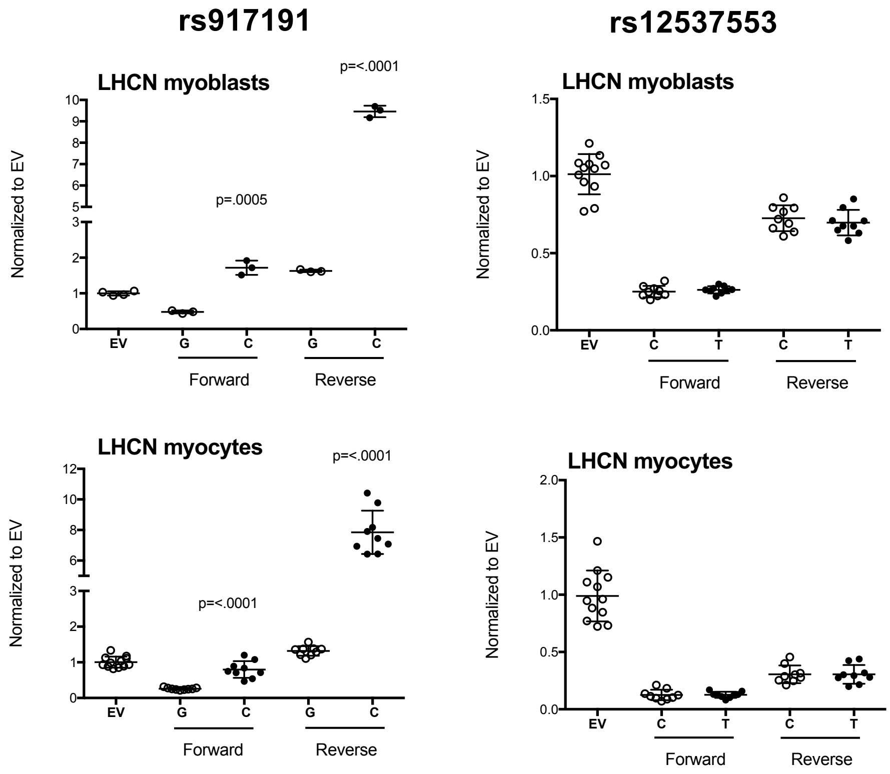


**Figure S16. Transcriptional reporter assays in myoblasts and myocytes for two WHRadjBMI candidate variants near *SEMA3C*.**
Transcriptional reporter assays testing function of rs917191 (left) and rs12537553 (right) alleles in LHCN-M2 myoblasts (top) and LHCN-M2 differentiated myocytes (bottom). Points indicate relative luciferase activity of independent clones compared to empty vector and lines represent standard error bars. *P*-values are for Student's unpaired t-tests to compare alleles. rs917191 shows strong effects and significant allelic differences in transcriptional activity.
